# Long-term quality assessment and monitoring of light microscope performance through accessible and reliable protocols, tools and metrics

**DOI:** 10.1101/2021.06.16.448633

**Authors:** Orestis Faklaris, Leslie Bancel-Vallée, Aurélien Dauphin, Baptiste Monterroso, Perrine Frère, David Geny, Tudor Manoliu, Sylvain de Rossi, Fabrice P. Cordelières, Damien Schapman, Roland Nitschke, Julien Cau, Thomas Guilbert

## Abstract

Reliable, reproducible and comparable results are what biology requires from microscopy. To achieve that level of confidence, monitoring the stability of the microscope performance over time with standardized quality testing routines is essential for mining quantitative data. Three levels of microscope quality control procedures should be considered: i) usage of accessible and affordable tools and samples, ii) execution of easy and fast, preferably automatized, acquisition protocols, iii) analysis of data in the most automated way possible with adequate metrics for long-term monitoring. In this paper, we test the acquisition protocols on the mainly used microscope techniques (wide-field, spinning disk and confocal microscopy) with simple quality control tools. Seven protocols specify metrics on measuring the lateral and axial resolution (Point-Spread Function) of the system, field flatness, chromatic aberrations and co-registration, illumination power monitoring and stability, stage drift and positioning repeatability and finally temporal and spatial noise sources of camera detectors. We designed an ImageJ/FiJi java plugin named MetroloJ_QC to incorporate the identified metrics and automatize the data processing for the analysis. After processing and comparing the data of microscopes from more than ten imaging facilities, we test the robustness of the metrics and the protocols by determining experimental limit values. Our results give a first extensive characterization of the quality control procedures of a light microscope, with an automated data processing and experimental limit values that can be used by core facility staff and researchers to monitor the microscope performance over time.

## Introduction

Producing reliable and comparable results is essential for quantification analysis in biology using fluorescence microscopy. A reliable quantification can only be possible if the quality of the observations is trustworthy. Furthermore, recurrent quality control (QC) testing of the microscope would be necessary. To obtain consistent quality, the microscope should be stable over time and every deviation from the standard values should be understood and corrected if possible. Moreover, understanding the limitations and deviations of a microscope’s performance is a way to understand better the fundamentals of microscopy, what properties and performances can be expected from a specific setup and how to troubleshoot your system if necessary. It also helps to build up a vocabulary necessary to interact with the microscope and devices manufacturers.

Determining the QC procedures, i.e., tools, protocols and analysis methods, is crucial and will define the efficiency of the microscope performance monitoring. Readily available tools, fast and robust acquisition protocols and automatized analysis methods can make the monitoring easily accessible to a broad scientific community. This task, often underestimated and thus neglected, is necessary and is critical in Life Sciences to ensure reproducibility among laboratories and over time monitoring ^1–3^ .

In the past, different groups have developed measurement protocols and published methods for a variety of wide-field and confocal microscope performance aspects ^4–7^. The majority of these QC studies were recently reviewed ^8, 9^. Some of the studies were carried out at a more international scale involving a larger community in the frame of the Association of Biomolecular Resource Facilities (ABRF) ^10, 11^.

In a few of these studies limit values and automatization analysis workflows were presented. An ISO norm for confocal microscopy was recently published ^12, 13^, providing a fixed minimal set of tests to be performed. This norm is descriptive, no experimental values are shown, nor are limiting values proposed, but it is a first step towards quality control standardization in microscopy.

In our work we provide experimentally tested and validated QC guidelines. For the first time we collected experimental data from over ten light microscopy core facilities and automatized the acquisition and analysis procedures based on commonly accepted protocols. The involved facilities are part of the French microscopy technological network RTmfm and this study results from of a collective work of the QC working group (GT3M) of this network.

Our intent here is to provide detailed open source protocols, affordable hardware and software tools and define reference values to assess fluorescence microscope QC metrics.

We focus on the following quality assessments:

i. Lateral and Axial resolution of the microscope. Images from single point emitters (e.g., single fluorophores in a sample) do not appear as infinite small points on a detector of a conventional microscope but as wavelength-dependent size-limited spots with an extension in order of the wavelength of the excited and emitted light; they are diffraction-limited die to the diffraction of light within the optical system ^14–17^. The size, shape and symmetry of the Point Spread Function (PSF) are characterized by the objective lens and other lenses used and the total optical beam path. Due to technical limitations the PSF is rarely close to the theoretical values given by optics, and thus the recorded image resolution is rarely as good as the expected one. Therefore, the image quality and the subsequent quantification analysis are affected. Evaluating and monitoring the PSF of a microscope over time is the first key step to determine the performance stability of a microscope and was well studied in the past ^17–20^ even with dedicated software tools ^21, 22^. This study validates our acquisition and analysis tools and protocols, and determines some experimental tolerance values.
ii. Field illumination. Imaging of a uniform fluorescent sample can be used to test the field uniformity of the system’s field of view. Ideally, all pixels should have the same value in this configuration. Unfortunately, illumination light source alignment and optical aberrations (from the objective lens and additionally optics included in the light path) can affect the homogeneity of the field illumination. Whereas, laser and LED light sources are coupled to the microscope with optical fibers and relay optics that help homogenize the beam’s Gaussian profile, not perfectly calibrated scanners can influence the laser illumination uniformity as well as the detection uniformity of laser scanning confocal microscopes. Bulb-type sources like metal-halide, mercury or xenon arc lamps are not very homogeneous to begin with and additional optical elements like diffusers are often necessary. Furthermore, the dichroic mirror and the filter positioning in the filter cubes can impact the observed field illumination for every color channel. Accurate image data quantification requires characterizing the field illumination pattern and correcting for heterogeneities if necessary. It is essential for image stitching, ratiometric imaging, tracking, segmentation experiments and even Fluorescence Resonance Energy Transfer (FRET) applications. When non-uniform illumination occurs in experiments involving tile scans (acquisition of adjacent XY planes), it leads to optical vignetting and unwanted repetitive pattern in the reconstructed image^23^. An image of the non-uniformity, as a reference, is then a prerequisite to correct this shading effect. Previous studies investigate the field illumination measurements by defining theoretical metrics and measurement protocols ^5, 11, 24, 25^. Here we propose three metrics to fast and easily characterize the field illumination using simple tools. These metrics are tested experimentally and we define some limit values. Two out of the three metrics are similar to the metrics defined by the ISO 21073:2019 norm for confocal microscopy^13^.
iii. Co-registration in xy and z. When the specimens are labelled with different colored fluorophores, a wrong alignment of the excitation sources (if several lasers or LEDs are used), insufficient chromatic aberrations corrected optical elements, mis-aligned filter cubes/filters, the use of non-pixel shift free multi-wavelength dichroic mirrors, or misaligned camera(s) will massively impact the resulting overlay image. To evaluate the co-registration/co-alignment, we used multi-labeled microspheres (i.e., beads incorporating multiple colored dyes) ^11, 19, 26^. Under ideal conditions, one would expect an excellent co-localization for all the acquired colors with these microspheres. The co-registration measurement will characterize the chromatic shifts of our system. These values can be further used to correct the acquired dataset (e.g. for colocalization studies, ratiometric or FRET imaging). However, accurate correction should be performed at the subpixel level and tackling the co-registration is more efficient at the hardware level than using postprocessing. A spatially resolved analysis of co-registration might also help to identify the region of interest of the field of view of the detector where default can be considered as negligible. We used the ratio of measured distance versus theoretical resolution as the metric to evaluate the co-registration quality of the system for each color pair. The shortest wavelength of the color pair was used to calculate the theoretical resolution, as it is the most stringent resolution value.
iv. Illumination power stability. Spatially resolved quantification of a fluorescent marker of a given cell component involves fluorescence quantification. Such emission quantification relies on the fluorochrome’s photophysical properties and is linearly related to the dye concentration and the intrinsic power of the light source used under conventional excitation. Accurate image quantification is thus essential and excitation and emission collection should be kept spatially uniform and constant over time to allow spatial and temporal comparison of fluorescence signals and, hence, dye concentration or labelling molecule density. Besides sample preparation issues and fluorochrome performance, excitation is the leading cause of variability, especially for researchers who want to compare samples acquired weeks apart. We propose to monitor light source fluctuations at three different time scales, long, mid and short-term. Compared to past studies ^11, 27^, we define the limit values of the metrics based on experimental values of a wide range of microscopes. We focus our study on laser sources for wide-field (TIRF), spinning disk and confocal microscopy. The long-term scale monitoring evaluates the illumination intensities of the sources over months or years and to what extend different experiments carried out on different days can be compared for quantification. Mid-term scale will provide crucial information on the fluctuations of the source intensity for typical live-cell experiment durations (e.g., overnight experiment, where room temperature stability is also crucial to ensure the illumination stability during the entire experiment). Short-term scale monitoring highlights potential variations during fast time-lapse acquisitions. This can either allow correction of experimental variations (provided monitoring is performed during the acquisition) or give confidence in fluorescence quantification accuracy, when fluctuations are negligible.
v. Stage drift and positioning repeatability. The stage stability is of great importance when performing time-lapse or multi-position imaging. The stability can be inherent to the stage characteristics or most often influenced by external parameters. Very often during time-lapse imaging in live-cell experiments or recordings of large z-stacks (hundreds of planes) instabilities can occur, which cause a pseudo movement of the observed structures over time. The setup can be affected by lateral and axial drift (change of the defined focal observation plane). Axial drift is a typical problem when using high NA objectives with a shallow depth of field. The causes are complex and can be due to thermal drift (temperature variations and thermal expansion), a thermal gradient in a temperature control chamber, mechanical movements (this could be the stage itself, the sample holder or the sample in the holder) or vibrations^28^ . Using home-made bead slides, we performed long period time-lapse experiments (duration of 15h) and characterized the 3D drift of various microscope setups. We characterized a stabilization time and calculated the 3D drift rate before and after stabilization. We experimentally defined tolerances values for these three metrics. For multi-position imaging the positioning repeatability of the stage should be high enough to image the same fluorescent points in the field of view (FOV). The repeatability is a specification of the stage and can be experimentally measured. We define repeatability as a measure of the ability of the stage to reposition itself to a reference point after moving in both the x and y directions. The repeatability error should stay constant during the lifetime of the microscope stage. Protocols exist in literature about repeatability measurements of stages. Unfortunately, the stage performance is often considered isolated ^29, 30^, and not as a part of a whole system. We considered the stage being part of the microscope system and defined a metric easily applicable. We compared the acquired experimental values to the stage specifications found in the manufacturer’s datasheets.
vi. Detector noise The last element in the acquisition chain of a light microscope is the detector. It can be a two-dimension (2D) array detector (camera type: CCD, EMCCD, sCMOS, others) or a point detector (PMT, APD, HyD, GaAsP, others). At low light levels, the detector noise can exceed the fluorescence signal to be detected. Therefore, one needs to quantify its contribution. But noise should not be confused with the background signal that depends on the sample itself. In the view of the different detector technologies, it is particularly complex to assess a single protocol that would apply to all cases. First, one should decide what to characterize, the detector itself or the whole detection system, considering the detector as the last element of a chain, where all previous components influence the captured signal. Previous QC microscopy studies mainly refer to the latter definition and evaluate noise within the framework of a complete detection system^5, 27, 31^. Although this method is beneficial and necessary for microscopy experiments when using actual samples, it is very dependent on any microscope components. Their influence of the upstream elements may hinder the ability to characterize the detector accurately. In our study we evaluated the noise of the detector itself to have some easily reproducible values. The noise sources of a detector can vary, although the main ones are those linked to fluctuations of thermal generation of electrons, due to readout noise, dark noise or due to electron multiplication (PMT, APD, EMCCD). The European Machine Vision Association has published standards on noise characterization of cameras ^32^. We propose here an easy and straightforward protocol to globally measure some camera’s noises. Read noise is the noise that s created during the conversion of photons to electrons and originates mainly from on chip amplifiers. It is measured in dark conditions to avoid shot noise contributions due to uncertainties of the photon arrival. Read noise is often a technical bottleneck when a low signal is to be quantified. This detector parameter is always provided in the manufacturer’s datasheet. Quantitative image data analysis, especially for maximum likelihood estimation methods, requires individual pixel characteristics, such as variation of the read noise ^33, 34^. We propose a protocol to quantify the read noise of CCD, EMCCD and sCMOS cameras on the installation day and later on. In that same protocol, the dark offset signal value is measured (a value arbitrarily adjusted by the manufacturer to allow access of the complete read noise), together with the dark signal non-uniformity (DSNU), which is a spatial noise. The DSNU refers to the non-uniformity between pixels, is measured in electrons and is the standard deviation of all pixels dark offset. In sCMOS image sensors DSNU is higher than in single output CCDs due to the pixel transistors and the column amplifiers of the sensor. EMCCDs often have additional noise distributed over the image sensor area. Compared to previous read noise and DSNU studies ^35^, we performed measurements among a wide variety of CCD, EMCCD and mainly sCMOS cameras for which in-depth noise characterization is missing since this kind of cameras are quite recent. We also provide an analysis tool and propose additional experimental tolerance values.

### Data analysis

To analyze the data, we developed an ImageJ/Fiji MetroloJ_QC plugin that fully automatizes the QC data analysis, based on existing MetroloJ plugin tools^36^. Processing was automatized to minimize user actions. We also implemented multichannel image processing to avoid splitting channels and separately configuring the plugin for each channel. The plugin also incorporates new metrics defined in that study and new tools for camera characterization and drift monitoring. It allows the creation of report documents to quickly identify parameters within/outside tolerances. The metrics were validated experimentally through a series of tests for many different manufacturer microscopes among more than 25 microscopes of ten core facilities using the same QC tools and protocols. Most of the microscope manufacturers are represented, with the majority coming from Carl Zeiss Microscopy (50%), followed by Leica Microsystems (20%), Nikon (14%), Olympus (9%). The remaining systems were microscopes from Andor and Till Photonics.

Although some company staff provide some QC test results upon installation, they are either incomplete or the tools used are not available for customers. As on-site installation measurements are mostly not systematically available, it is currently up to the customer to set up protocols to carry out first (and subsequent) measurements and have initial values for further comparisons. Here we suggest that these first acquisitions and measurements could be an excellent way for users to be trained by manufacturers to correctly use a microscope. Then, the quality control monitoring over time can start, using tolerance values that we provide here as the first reference. With time and experience, the users may set their experimental limits, so that they can use to highlight issues and undertake an action or a service visit. Interestingly, limit values are required for quality control procedures, which provide valuable indicators to improve the service and the overall quality standard of a core facility.

## Materials and Methods

### Point Spread Function

#### Samples

We used sub-resolution beads for measuring the PSF of the high numerical aperture (NA) lenses. In the same slide we also mixed larger beads (1 μm and 4 μm) for easy detection of the focus plane and co-registration and repositioning measurements. The aim was to have one simple, multi-size bead, cheap slide that serves for many metric measurements.

We used 175 nm diameter blue and green microspheres for the PSF measurements, for DAPI and GFP channel respectively (beads PS-Speck, #7220 Thermo Fisher Scientific). Beads were vortexed to avoid aggregates. 50 μl of the diluted solution was dried overnight at room temperature in the dark on ethanol cleaned Zeiss 18 mm #1.5 High-performance square coverslips. The desired bead density on the slide is 10 beads in a 100x100 μm field. The coverslips were mounted on slides with 10 μl of ProLong Gold antifade mounting medium (#P36930, Thermo Fisher Scientific, refractive index 1.46).

As beads are directly next to the coverslip, any spherical aberration/distortion, induced by the potential refractive index mismatch between mounting medium and lens immersion medium are minimized. Other configurations may be used to control the depth of the beads within the mounting medium and monitor the effect of the RI mismatch on image quality (as to reproduce real in-depth conditions of a typical biological sample). The slides were left for three days at room temperature in the dark, to let the mounting medium fully cure. After three days, the samples were sealed either with Picodent dental silicone (Picodent twinsil®, picodent, Dental Produktions und Vertriebs - GmbH) or with nail polish.

To compare the effects of the bead size on PSF measurement results, we used yellow 100 nm beads (FluoSpheres, #F8803 Thermo Fisher Scientific), mounted as previously with the same preparation method.

#### Acquisition protocol

For PSF measurements, immersion objectives were tested and some 20x dry objectives with numerical aperture higher than 0.7. For the blue channel, the beads were imaged on wide-field microscopes with settings used for DAPI imaging (typically a bandpass 350/50nm excitation filter, a 400 low pass dichroic beamsplitter, and a 460/50 nm emission filter). On laser scanning confocal microscopes (LSCM) the beads were excited with a 405 nm laser line and fluorescence was collected between 430-480 nm. For the green channel, the beads were imaged with the wide-field microscopes with similar setups like for GFP imaging (typically with 470/40nm and 525/50nm bandpass excitation and emission filters with a 495 long pass beamsplitter) and for LSCM beads were excited with a 488 nm laser and fluorescence was collected between 500-550 nm.

The optical sampling rate is crucial for accurate PSF measurements. At least the Shannon-Nyquist criterion should be fulfilled. Most of the collected was even slightly oversampled, in order to get a more precise fitting. Some exceptions were the cases of the lateral X/Y sampling and the 20x dry objectives with NA > 0.7 coupled with cameras used in classical wide-field setup, lacking any other magnification relay lens. Whereas the typical theoretical lateral resolution around 350 nm for GFP channel, the typical minimum pixel size is around 6.5 μm in a wide-field setup, corresponding to 325 nm of pixel size on the image. For the same reason, respecting the Shannon-Nyquist with EM-CCD cameras having a physical pixel size of 13-16 μm was difficult in some configurations.

Concerning the lateral Z sampling, the Shannon-Nyquist criterion was also considered which gave for instance 0.15 and 0.2 µm step size for the highest NA objectives (1.4) for confocal and wide-field microscopy respectively.

For LSCM imaging the confocal pinhole was set to 1 Airy unit (AU) to assess the standard performance of a confocal microscope, keeping a high ratio of signal intensity to resolution. In the case of a weird PSF (not straight or not symmetric along the z axis for instance), the pinhole was widely opened to record the entire aberrations. Very often, pinhole alignment was necessary before or after the tests. Moreover, as shot noise affects the PSF measurement, a high signal-to-noise ratio is essential for the analysis fitting and the precise Full Width Half Maximum (FWHM) calculation. We found that single color beads, compared to multi-color ones, provide a higher signal and should be used preferably. A tutorial video summarizing all these steps was produced by the GT3M of the RTmfm network (https://youtu.be/ll4X_e8_mo8).

#### Metrics

The experimental PSF values were calculated using the MetroloJ_QC plugin with the Fiji software. The plugin calculates FWHM along the x, y and z axis of all beads in the field of view. Before analyzing the images with the plugin, we visually inspected the acquisitions. Some images needed to be cropped to remove saturated beads. Alternative options can be used to discard any saturated beads from image. Only beads that were close to the center of the FOV (30% of the entire sensor for sCMOS, zoom of 6 for confocal) were analyzed to avoid aberrations. At least five beads were analyzed.

The MetroloJ_QC plugin works as follows: First, the xy coordinates of beads are identified using a find maxima algorithm on a maximum intensity projection of the stack. Beads too close either to the image’s edge or to another bead are discarded from the analysis (the user in the plugin can modify this distance). Then, FWHM is measured: the maximum intensity pixel within the entire 3D data set is determined and an intensity plot along a straight line through this maximum intensity pixel point is extracted in all dimensions. Finally, the bell-like plots/curves are fitted to a Gaussian function using the built-in ImageJ curve fitting algorithm to determine the FWHM.

The experimental FWHM was compared with the following theoretical resolutions:

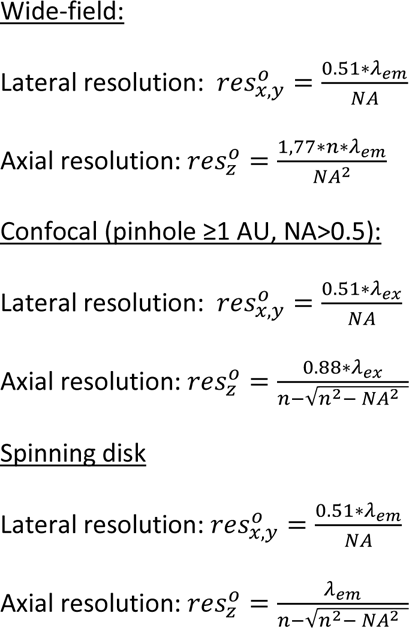

Where λ_em_ is the emission wavelength, λ_ex_ the excitation wavelength, n is the refractive index of the immersion liquid. Resolution formulas are the FWHM of the PSF and not the distance of maximum to the first minimum of the intensity profile of the PSF ^37, 38^. For spinning disk confocal microscopes, we consider that the pinhole size does not fill 1 AU (it is the case when we use a circular pinhole of size close to 50 μm) ^39^.

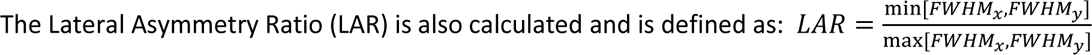

#### Analysis

After automation of the FWHM measurements on several beads/datasets, average values are extracted in all three dimensions, and standard deviations are calculated. Aberrant “PSF” (non-bead features that are considered as beads using the algorithms) can be further filtered using the Gaussian fitting parameter R^2^ (square R stands for the coefficient of determination, showing the goodness of a fit of simulated vs measured values) for each FWHM measurement. We recommend a value superior to 0.95. Further inspection of each PSF, using the computed bead signal to background ratio, will help removing aberrant values. Additional parameters are also measured, like the lateral symmetry (named asymmetry parameter), or whether the acquisition meets the Shannon-Nyquist criterion.

The figures in that manuscript and the calculations on the statistical significance are prepared with the GraphPad Prism software.

### Field illumination

#### Samples

For the field illumination flatness assessment, fluorescent plastic slides can be used (blue, green, orange and red, provided by Chroma Technology Group #92001 or Thorlabs #FSK5). Those slides are very bright. Thus we used a not optimally excited slide at the tested wavelength (for example the orange or red slide for a 488 nm excitation). Their inconvenient is their low signal in the far-red channel (>700 nm emission). For a complete field illumination characterization, one should perform measurements for all four main channels (DAPI channel to Cy5 channel). For a more rapid and regular characterization, a field illumination measurement at the DAPI and GFP channels can be enough. A glass coverslip can be sealed on the plastic slide with immersion oil to protect the plastic slide from scratches ^27^. We compared the plastic slides with a home-made homogeneous thin fluorescent specimen. We used a highly concentrated (1mM) dye solution (rhodamine) between a glass slide and coverslip for this specimen. A low-cost alternative that seems to work efficiently is a thin layer of a fluorescent marker pen (e.g., Stabilo) on a coverslip. We performed some comparison tests of the three above samples.

#### Acquisition protocol

For wide-field and spinning disk microscopy the whole camera chip was used. For confocal microscopy we used the recommended zoom of the manufacturer to yield a uniform field of view (usually it is a minimal zoom value of 1), as suggested in the ISO 21073 :2019 norm ^13^. We placed the slide on the stage and focused on the surface of the slide. We measured the field illumination slightly deeper than the surface level (around 5-10 μm) to avoid recording scratches or dust. The acquisition parameters were set to take full advantage of the detector dynamic range and avoid saturation.

#### Metrics

We defined three metrics to evaluate the field illumination:

i. The uniformity of the illumination at the observed FOV, U[%]. It is expressed as follows: 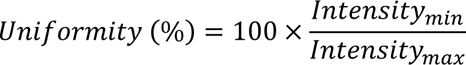

where Intensity_max_ and Intensity_min_ are the maximum and minimum intensity acquired in the field of view, respectively. The size of the field of view influences the value of this metric.
ii. The field Uniformity (fU). It is the standard deviation of the normalized intensity relative to the maximum intensity of the FOV at the four corners and the four middle-points of the four sides (where absolute intensity is normalized between 0, corresponding to the minimum intensity in the image and 1, corresponding to the maximum intensity found in the image) and is expressed as follows:

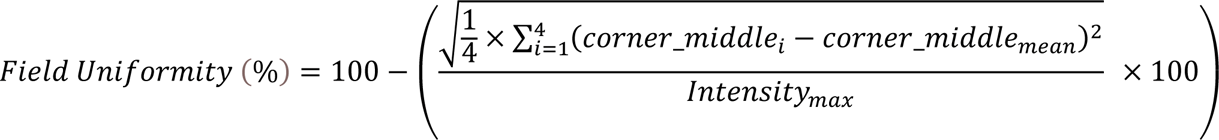
iii. The centering of the maximal zone of the intensity of the image, C[%]. It is expressed as follows:

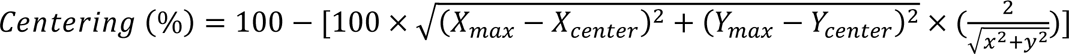

where X_max_ and Y_max_ are the coordinates of the “maximum intensity” zone, X_center_ and Y_center_ the coordinates of the center of the FOV, x and y determine the width and the height of the image. The size of the field of view does not influence the metric value (normalization using the x and y values of the image). The Uniformity and the Centering metrics are included in the ISO 21073 :2019 norm^13^ and setup specifications using this norm should provide these values.

#### Analysis

The MetroloJ_QC plugin locates both minimum and maximum intensities within each channel. It also finds the center of the intensity. As in the MetroloJ plugin, a normalized intensity image is calculated. Then, using the normalized image, the “maximum intensity” location is determined considering a reference zone (either the 100% intensity, i.e. the geometrical center of all pixels with a normalized intensity of 1, or any other zone, such as 90%-100%, i.e., the geometrical center of all pixels with a normalized intensity between 0.9 and 1).

The user is prompted to divide the normalized image intensities into categories (bins). This value will be used for the computation of the reference zone and generation of the iso-intensity map. A value of 10 will generate iso-intensity steps of 10%. A threshold is then used to locate all pixels with the maximum 100% intensity or the last bin window. The geometrical center of this reference zone is then located. Distances of these points to the geometrical centre (Center of intensity, maximum intensity, Center of the thresholded zone) are computed, and the above metrics are calculated.

The user can choose to discard or not a saturated image. Whenever saturation occurs in a few isolated pixels, noise may be removed using a Gaussian blur of sigma = 2. Note that saturation computation is done after the Gaussian blur step. Hence, whenever aberrant saturated isolated pixels are polluting the channel, if Gaussian blur gets rid of them, the image will no more be considered as saturated as no saturated pixels will be found. In the case of “clusters” of saturated pixels, the applied Gaussian Blur is not strong enough to eliminate the cluster center’s saturated pixels and the channel will still be reputed as saturated and skipped if the discard saturated sample option is selected.

A batch mode enables the automatization of the analysis.

### Co-registration

#### Sample

For the co-registration measurements, a home-made bead slide containing 1 μm and 4 μm four color beads is used. The 1 μm beads (#T7282, TetraSpeck, Thermo Fisher Scientific) had a final density on the slide of around 1bead/100μm^2^. The 4 μm beads (#T7283, TetraSpeck, Thermo Fisher Scientific) were more diluted to achieve a density of 1 bead/300μm^2^.

#### Acquisition protocol

3D stacks of multicolor fluorescent beads in two or more channels were collected along at least 10 μm. For LSCM systems the zoom was higher than 3. The Shannon-Nyquist criterion was met whenever achievable and saturation was avoided. At least five beads were acquired, coming from at least two different FOV. The beads should be very close to the center of the image (no more than one fourth of the total field of view) since objectives are usually best corrected for chromatic shift in their central field. Alternatively, some characterization of evenness of the chromatic aberration can be performed, using different beads locations within the field of view.

#### Metrics

Our metric here is the r_experimental_/r_reference_ ratio. In more detail, the acquisition consists of 3D stacks with two or more emission channels. MetroloJ_QC uses automation of the elegant MetroloJ coalignement algorithm. Beads are identified, saturated beads or beads too close from the edges/another bead are discarded. Then the center of mass for each channel is calculated and the distances between the centers are estimated (r_exp_). For each channel to channel distance r_exp_ a reference distance is calculated (r_ref_) by taking into account the resolution in that position and angle at the highest emission wavelength ^40^. The plugin calculates the ratio r_exp_/r_ref_ for each color couple. We then quantify the number of cases that have a ratio below or above 1 for each color couple and each technique.

#### Analysis

The image analysis was performed using MetroloJ_QC. It automatically generates analyses for all channel combinations possible, measures the pixel shifts, the (calibrated/uncalibrated) intercenter distances, and compares it to their respective reference distance r_ref_. A ratio of the measured intercenter distance to the reference distance is also calculated. Images with more than one bead can be analyzed and a batch mode enables analyses of multiple datasets.

### Illumination power stability

#### Sample

Different strategies for the evaluation of illumination power stability exist. We used the power-meter console (PM100D, Thorlabs) coupled to a microscope slide power photodiode Si sensor, 350-1100 nm sensitive, designed to measure optical powers from 10nW – 150mW (S170C, Thorlabs) with a response time of 1 μs. This sensor fits in the microscope slide holders of upright and inverted microscopes and measures the power at the sample plane. The controller console is connected to the computer by USB. The illumination power can be recorded over time with the Power Monitor GUI software at a defined sample rate. When needed, we monitored the temperature with the TSP01 temperature probe (Thorlabs) and recorded it with the TSP01 Application software.

#### Acquisition protocol

We chose a low magnification objective (dry 10x, NA 0.3 or 0.4). We clean the objective before use, in case of any remaining dried dirt (oil, glycerine) on the lens. The slide power sensor was placed on the stage and the objective was centered on it. We adjusted the power-meter wavelength correction to the corresponding laser wavelength we wanted to measure. The background correction was performed in the dark. The laser should be warmed up and switched on at least one hour before the measurements. We regulate the laser power at 100% of the maximum power, eliminate laser noise and ensure that the laser is in its stable working range. In some cases, the 100% power can damage the power meter sensor if the measurement is done in a continuous mode (often for TIRF of SD microscopes), so in these cases we decrease the power at 20%. For confocal microscopy, we defined a bleach point in the center of the FOV (laser beam stationary) and launched a time series. If no bleach point mode is available, the highest zoom was applied at bidirectional mode. We prefer the bleach point mode to avoid AOTF modulation of scanning laser beams. The blanking function was switched off for TIRF or spinning disk microscopes.

#### Metrics

We divided the stability metrics into three time scales: short, mid, and long time scale. For short-time scale stability, the power was measured every second and recorded for a duration of 5 min, after having turned on the source for at least one hour. For mid-time scale stability we measured values every 30 sec for a duration of 2h. The source was constantly ON, illuminating the detector (eg. laser at bleach mode with AOTF ON), during the measurement period. For these two time scales we calculate the standard deviation (STD) of the measurements and the stability factor (STAB).

The stability is defined as:

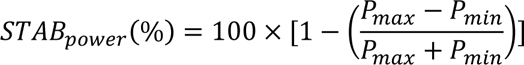

with P_max_ and P_min_ the maximum and minimum recorded power. The stability factor is the one defined in the ISO 21073:2019 norm^13^. For long-term monitoring we measured every month the laser power and traced the evolution of the source power through its lifetime.

#### Analysis

We calculate the STD and stability factor for the short and mid-time scale stability measurements. For the long-term scale stability we trace the values over the laser lifetime. We do not calculate the STD and STAB, as their values depend highly on other parameters than the laser stability itself, as its misalignment in time or the aging of the laser.

### Stage Drift

#### Sample

The home-made fluorescent bead slide with the 1 μm diameter beads or the 4 μm beads was used (same as for co-registration studies).

#### Acquisition protocol

We placed the sample on the stage that was switched on at least 30 min before the experiment, like the whole microscope setup and its components. We centered one bead in the illumination field and adjusted the z-position on focus. We started z-stack recordings over time on a range of 15 μm. We preferred using a 60/63x high NA oil immersion objective, to be close to the experimental conditions when overnight experiments are started and measured precise XYZ variations. We evaluated the long term drift (overnight acquisitions) for a time-lapse of 15h with an acquisition frequency every 10 min at room temperature. In the microscope rooms the temperature is usually set around 21°C. However, variations of temperature lower than one degree were possible, but were not monitored.

#### Metrics

We measured the mean velocity of the beads in μm/min, the stabilization time of the stage and the subsequent velocity after this time. The stabilization time is the time during which any axis displacement within 10 minutes should be kept below a high NA minimal resolution distance of 200 nm, so that drift should be barely noticed during 10 minutes in diffraction-limited conditions. This is considered as a mild drift if all three axis displacement speeds are less than 15 nm/min for each axis, a drift that can be neglected in diffraction limited experiments.

#### Analysis

We used the TrackMate plugin^41^ of Fiji which is widely used for single particle tracking purposes. The plugin finds the center of mass of the bead for each time-point in the three dimensions and it calculates the mean velocity of the bead and the 3D displacement. We exported the bead xyz displacement and traced it over time, to find the stabilization time. Finally, we recalculated the mean velocity for the time values after the bead stabilization.

### Stage positioning repeatability

#### Sample

All kind of samples that can be well-distinguished in fluorescence or transmitted mode would work for these kind of measurements. In our cases we use the 1 or 4 μm fluorescent multicolor beads of the self-made bead slide (same slide as for the coregistration studies).

#### Acquisition protocol

To determine the repositioning of the x-y stage movement we had to move between multiple fixed points over time and observe the variability of the position precision. First, the stage is initialized (whenever this option applies). The microscope with all its components is switched ON at least 30 min before the experiment. 10x or 20x dry objectives were used to avoid any oil pressure/surface tension influence. A bead slide is firmly fixed on the stage (usually the stages offer a clipping mode that fixes the slides). To avoid drift due to heating/cooling of the slide, the setup stays in this state for 15 minutes. A stage position, where a fluorescent bead sits in the center of the field of view, is recorded (referred as bead position P_ref_). The stage is then shifted 3 mm away in both x and y-axis towards the top-right direction (i.e., a maximum of 4.2 mm diagonal movement). This position (referred as top/right position P1) is recorded. Back to the bead position, the stage is further shifted away at the opposite position along the x-axis (bottom/right, 4.2 mm away from the bead position, referred as P2). The opposite positions along the y-axis are also recorded (referred as P3 and P4), resulting in a total diagonal movement of 8.4 mm. Then the acquisition software of the microscope is set to acquire a 20 cycles time-lapse of the nine positions (P_ref_-P1-P_ref_-P2-P_ref_-P3-P_ref_-P4-P_ref_ …). The above diagonal movements are the displacements that could result in the worst repositioning since they involve movement in both x and y axes.

A 4.2 mm distance was chosen as most often imaging of a coverslip/slide mounted sample does not involve more than 3 to 5 mm stage movements. This value has to be adapted depending on the application the setup is intended for. To fully characterize a stage for all kinds of applications, we perform repeatability measurements for two more distances along the x and y axis, for 0.3 mm and 30 mm when achievable. The latter distance corresponding to experiments on histological slides or multi-well plates.

For both stage drift and repeatability acquisitions the flatness of the stages was checked. Metrics

The x and y coordinates of the bead reference position are calculated for each cycle. Standard deviations of this coordinate are then compared to the repeatability values given by the manufacturer.

#### Analysis

Bead position is analyzed using the standard Fiji tracking plugin (e.g., TrackMate). We recorded the x-y positions in μm calculated for each cycle and then we calculated the standard deviations.

### Camera noise measurements

#### Sample

No sample is needed for this measurement. Acquisition protocol

This protocol was performed for CCD, EMCCD and sCMOS cameras. The CCD cameras were all Coolsnap HQ2 (Photometrics). The EMCCD cameras were iXON 888 (Andor) and Evolve512 (Photometrics). The sCMOS cameras were Zyla 4.2 or 5.5 (Andor), Orca Flash 4 LT, LT+, V2, V3, Fusion BT (Hamamatsu), Prime95B (Photometrics) and Edge 4.2 (PCO). When possible, camera shutter was closed to avoid any unwanted light contributions and thus unwanted shot and read noise. When available, the cameras were cooled according to the manufacturer recommendations (for sCMOS 0 or -10°C, EMCCDs -70 to -80 °C). The acquisition software was set to acquire a 100 cycle time-lapse. The detector exposure time was set to short values (e.g. 10 ms) to avoid Dark Signal Non Uniformity (DSNU) contribution and pixel binning was not used. This procedure is repeated using different camera settings (e.g., readout frequencies or gain values). For EMCCD cameras we set the EM gain at the minimal value of 1, which is not the common condition that these cameras are used but this setting allows comparison with the manufacturers’ values.

#### Analysis

The data analysis was performed using the MetroloJ_QC plugin. It generates both intensity average and standard deviation projections across the 100 cycles. The average intensity projection was used to calculate offset and DSNU values. The offset is the mean average intensity across all pixels of the average intensity projection. The DSNU is the standard deviation of the pixel intensities of the mean average intensity projection, multiplied by the electron to analog-to-digital units (ADU) conversion gain/factor. To find the read noise map, we calculated the standard deviation image of the 100 frames in electrons (the conversion factor e^-^/ADU should be used). The root mean square (rms) noise refers to the rms value of all pixels read noise. The median noise is the median value of all pixels read noise (dark noise is considered equal to zero for this short acquisition time).

We defined two metrics: the first consists of the variation in percentage of the measured read noise to the camera’s read noise as indicated in the datasheet (theoretical or the provided one for a specific camera). We define it as:

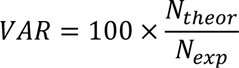

with N the read noise value (rms or median, depending on the given theoretical value) of the camera. The second one consists of the stability of the read noise over time and is defined as

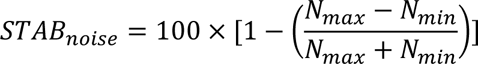

with N_max_ and N_min_ the maximal and minimal read noise values.

## Results

### Microscope Lateral and Axial Resolution (PSF)

In total, data from 91 objective lenses from more than 10 light microscopy core facilities was analyzed. Ten bead samples were prepared in the same way and distributed to the facilities. 63x/60x lenses were considered reference lenses and two-color PSF measurements were most often acquired (in the DAPI and GFP channels). For the other lenses, both colors were not always acquired because blue bead fluorescence is usually dimmer than green bead fluorescence and it was difficult to achieve good signal-to-noise ratio images when lower NA lenses were used.

We first investigated if the proposed procedure was robust enough (i.e., if the variability of the calculated PSF metrics with the same sample/setup was kept negligible). PSFs of the same bead at five different time-points were acquired (GFP channel). Figure 1a shows that for three different microscopes representing the wide-field, spinning-disk confocal or raster, single-point scanning techniques (WF, SD, LSCM) that we studied, the repeatability accuracy is quite high. The % variation (ratio of standard deviation to the FWHM) in x-y ranges goes from 1 to 5 %, while the z direction ranges from 4 to 6%. This shows that small variability exists among the acquisitions, it is higher for the z-axis (the Gaussian fitting is less precise than for x-y direction), but it rests quite low.

**Figure 1:**
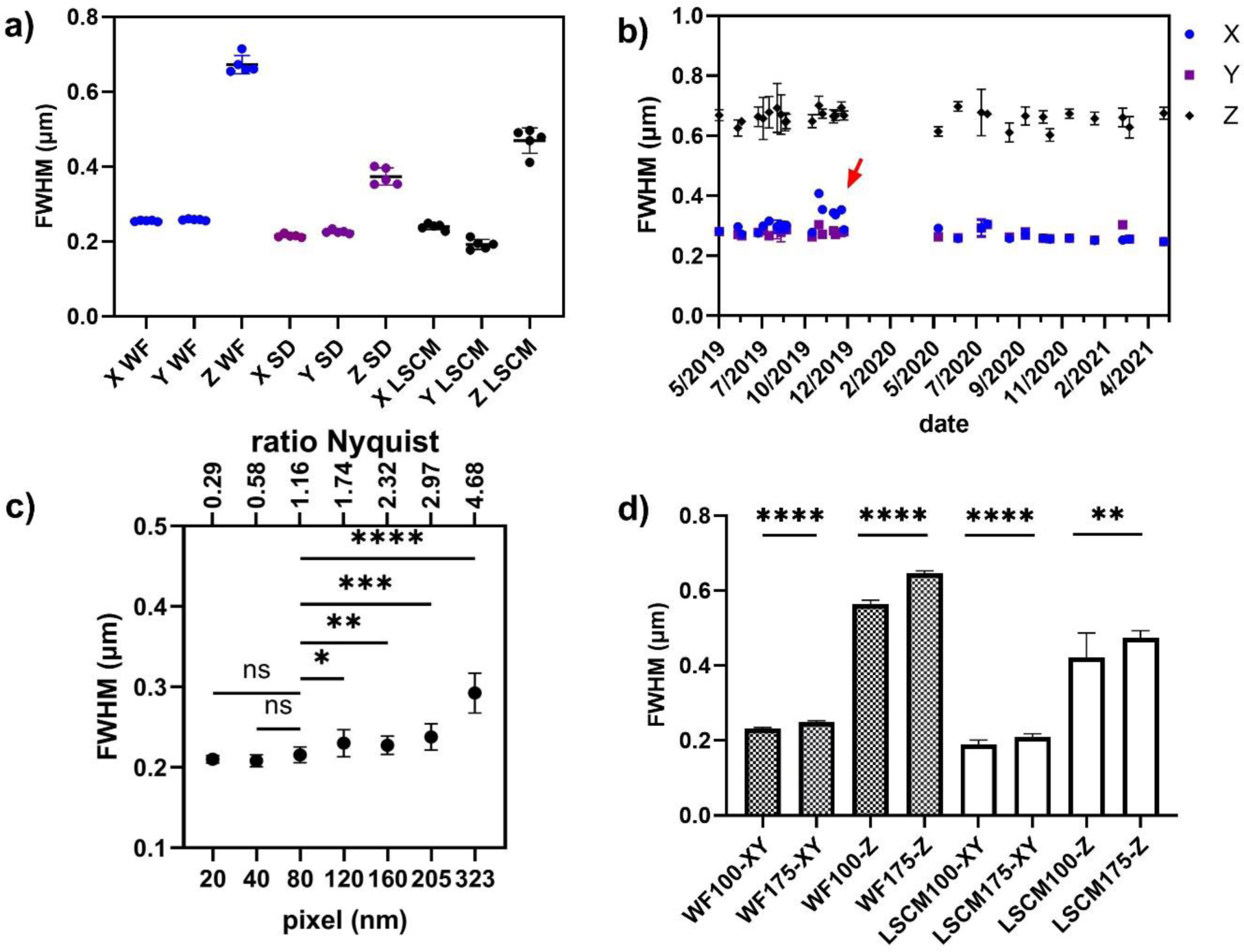
PSF variability measurements. a) PSF repeatability accuracy for wide-field, spinning disk and LSM microscopes along the x, y and z axis. PSF were acquired five consecutive times. b) Stability evolution of PSF over 22 months for an upright wide-field microscope. The red arrow shows when the PSF is significantly different along the x-axis. The gap at the dates of January 2020 corresponds to Covid lockdown; no experiments could be carried out. c) Lateral PSF dependence on the sampling rate on a LSCM microscope (1AU). Statistical significance was determined using Dunnett’s multicomparison test, with the 80 nm setting chosen as the control value. Two-tailed p-value test was nonsignificant (ns), <0.05 (*), <0.005 (**), <0.0005 (***) or <0.0001 (****). d) PSF dependence on microsphere size for wide-field (WF) and single point raster scanning LSCM microscopes. WF100 and WF-175 stand for wide-field PSFs for 100 nm and 175 nm beads respectively. Statistical significance was determined using t-test for each pair and each condition (one pair: 100 and 175nm beads). All measurements were done with a 63x lens, NA 1.4, at the GFP channel (525 nm emission).

We then measured the stability of the objective performances by analyzing PSFs acquired for over a year, using the same microscope, objective, sample and also acquired by the same operator (Figure 1b). Variations of lateral (x,y) and axial (z) FWHM were small (less than 5%). Regular analysis of objective performance is essential for facility managers. We also recommend facility staff to acquire PSFs (and other metrics) before and after any scheduled (e.g. annual) revision of a system, to compare QC data. In October 2019 (Figure 1b, red arrow) measurements show a 10 % increase of the x-axis FWHM, while no significant change is observed for the y axis FWHM. This variation follows an external service provider revision. After many tests, repositioning, and tilt adjustment of the camera, the symmetry was recovered (measurement of December 2019 and later on).

Another important consideration for PSF evaluation is the sampling density. One has to collect image with a high enough sampling density in the x-y and z dimension to get the most accurate PSF Gaussian fitting. The Shannon-Nyquist sampling density criterion was used as a reference. To experimentally determine the adequate sampling rate/density, high Signal to Background Ratio PSF acquisitions were performed using different voxel sizes. This analysis was done with a confocal setup since the pixel size can be easily changed (compared to wide-field setup, where, due to their fixed physical camera pixel size, the options, such as camera binning or use of a magnification relay lens, are limited). Figure 1c shows the calculated FWHM for different pixel sizes. The Shannon-Nyquist criterion was considered as met whenever pixel size is equal or lower to λ_ex_/8*NA ^42, 43^. Using a 1.4 NA lens and an excitation wavelength of 488 nm, the criterion value is 43 nm (Figure 1c). This is the case for a closed pinhole (near to 0.25 AU). For 1 A.U. pinhole, the Nyquist criterion allows a 1.6x bigger pixel size (https://svi.nl/NyquistRate). In our case, with a pinhole of 1 A.U., we observed that the pixel size has an influence on lateral FWHM values when the pixel size value is higher than a threshold of 80 nm or higher than 1.16x the Nyquist criterion value (statistically significant of p<0.05 when performing Dunnett’s multicomparison test, setting the 80nm pixel as the control).

A third consideration for accurate PSF measurements is the choice of the fluorescent microspheres used. Hence, their size and brightness can alter the FWHM calculation. The brightest beads should be used, and this is the reason we recommend using single labeled beads. The beads diameter should be below the resolution limit. Figure 1d shows measurements taken with two setups with a 1.4 NA lens. Using the wide-field setup and either 100 nm or 175 nm diameter beads, FWHM values were significantly different (either lateral x-y values or axial z FWHM, p values smaller than 0.05). With 100 nm beads, the FWHM values are very close to theoretical resolution. Using a raster scan single point confocal (LSCM), differences were observed for both lateral and axial directions, but along the z axis the differences were less significant. As 100 nm beads are dimmer than the 175 nm ones, we assume this decrease in significance is due to a lower Signal-to-Background Ratio with the 100 nm beads, degrading the axial PSF fitting accuracy. As much more photons get collected in the wide-field mode, this photon shot noise effect is not biasing the wide-field FWHM measurements.

We conclude that when looking for the most accurate PSF measurements, 100 nm beads should be recommended. However, for PSF monitoring over time, brighter beads are needed. They are also more convenient if a wide range of objectives is to be examined using the same slide. Hence, with 100 nm microspheres, dimmer than the 175 nm beads, and with low magnification/NA lenses, detecting the beads and achieving a correct FWHM estimation may prove quite challenging. We recommend using single color 175 nm diameter beads for reliable PSF monitoring for these reasons. A mix of different bead colors may be used to speed up image acquisition and enable, in a single z-stack, measuring FWHM for different wavelengths.

The last consideration was the signal to background ratio (SBR). We observed that if the SBR is low, then the precision on the FWHM calculation is low (Figure S1), thus paying attention to background is highly recommended.

Finally, we analyzed the results obtained with the three microscope techniques. Figure 2a shows the FWHM distribution for all tested objective lenses for DAPI and GFP channels. Obviously, for the DAPI channel the FWHM measures values are worse than those in the GFP channel, as the objectives and the optics of the whole system are set for the GFP channel. The adjustable pinhole of LSCM microscopes was set to 1 A.U. Attention was paid to the SBR in LSCM images. LSCM measured FWHM values stay close to the theoretical values (Figure 2b), showing a large dispersion along the z axis. Many cases that show a z value far from the theoretical values are due to spherical aberrations, especially when PSF is taken with water (blue arrows in Figure 2a) or air low NA (25x) objectives (black arrows in Figure 2a).

**Figure 2:**
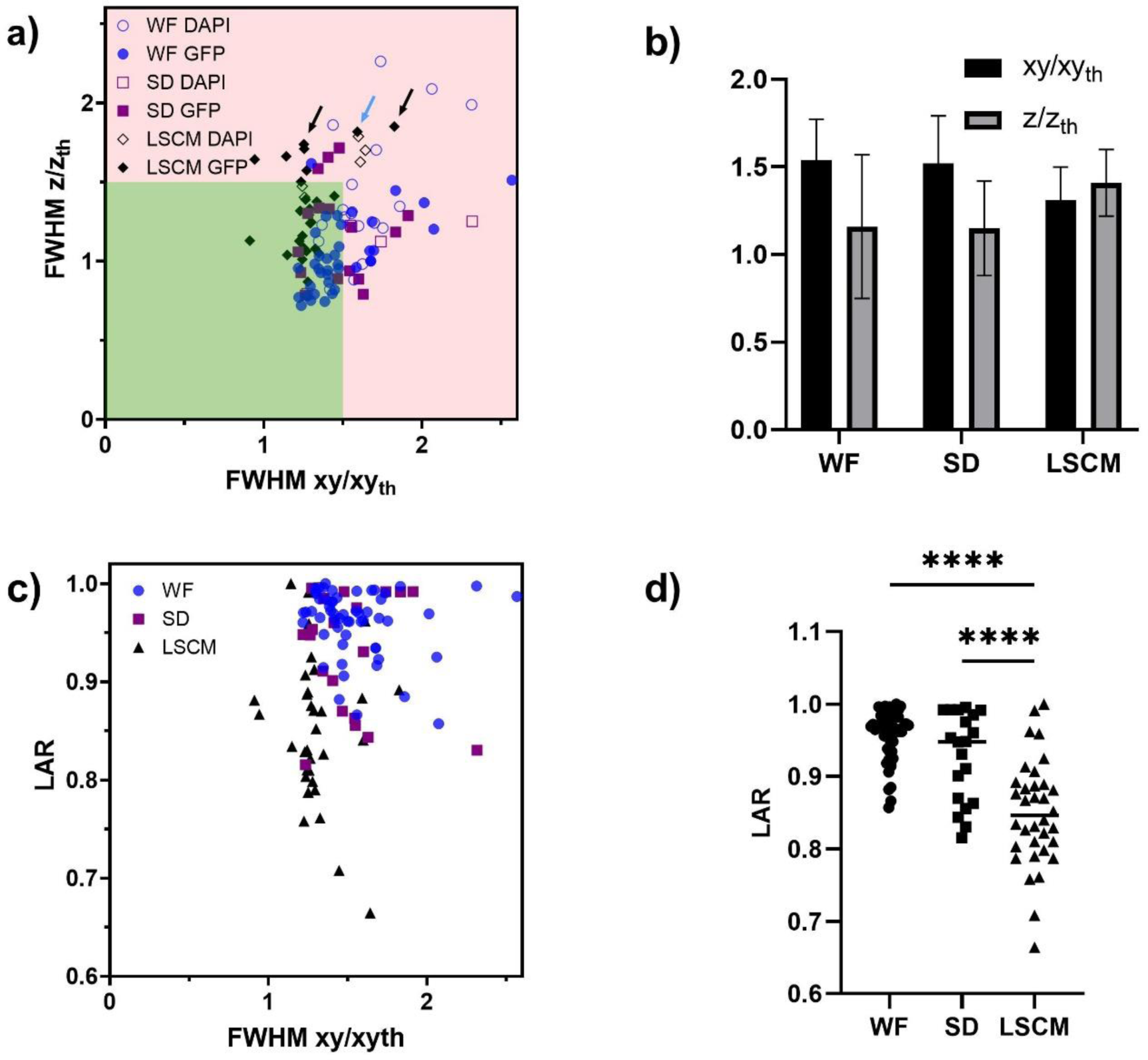
PSF distribution and asymmetry summary of wide-field, spinning disk, and LSCM microscopes. a) ratio of experimental to theoretical lateral and axial FWHM values for the three techniques and the DAPI and GFP channels. Black arrows show some dry 20x objectives, the blue arrow shows a water objective 40x, NA 1.2. b) Mean values of a). c) Lateral asymmetry of PSF. LAR is compared to the lateral FWHM experimental to theoretical ratio. d) LAR value for the three techniques and the statistics of the results (Two-tailed p-value was <0.0001 (****)). Statistical significance was determined by using a t-test.

The wide-field and spinning disk resolution measurements show similar behavior. The measured FWHM values are worse along the x-y axis than the z-axis, compared to the theoretical ones. For these two microscopy techniques, meeting the Shannon-Nyquist criterion is not an easy task (fixed physical pixel size of the detector array, not much further magnification possibilities). We believe under-sampling is the main reason explaining this discrepancy. Of note, as the sampling criterion was met in z, the observed ratios for the axial FWHM are closer to 1. For instance, a wide-field case that is far from theory is pointed out with a blue arrow (Figure 2a) and concerns 40x Plan APO 1.3 NA objective lens, with a pixel size of 183 nm.

The symmetry of the PSF is also an important parameter. Figure 2c shows the Lateral Asymmetry Ratio (LAR), defined as the ratio between the smallest and highest of both x and y experimental FWHM. We observed that for confocal microscopy the mean LAR is much lower than for the other two techniques (confocal: 0.85, spinning disk: 0.92, wide-field: 0.96). It is statistically significant as shown in Figure 2d (p-value for both comparisons <0.0001).

#### Field illumination

To check the field illumination for fluorescence microscopy, first, one has to pick the appropriate tools. The thin layer of a fluorescent dye and the fluorescently doped plastic slides are most widely used ^17^. The thin layer of a fluorescence dye gives a more precise illumination pattern and can be more convenient for shading corrections for tile scan acquisitions. The plastic slides are quite thick (some mm) and cause an index mismatch when used with immersion oil of the objective. For wide- field illumination or spinning disk microscopy, the collected background from the out-of-focus planes can alter the illumination pattern. For spinning disk, the pinhole crosstalk detection is well seen with plastic slides. An alternative and cheap solution for using thin dye layers and plastic slides is the usage Stabilo pens. We tested the three tools and found that for the defined metrics, the plastic slides are the most convenient (Figure S2).

We performed the flat illumination measurements for 130 objective lenses for a wide range of magnifications (5x-100x). Our reference channel is the GFP channel, although many of the measurements were performed for both the DAPI – GFP channel (alignment is often different between DAPI and visible channels) or for all the basic four channels. We defined three metrics, the first two are the Uniformity (U) and the Centering (C). U is the ratio between the minimum and maximal intensity values of the image. A ratio of 1 (100%) represents a most ideal case. C is the centering of the illumination in the image, normalized by the diagonal of the image. A value equal to 1 or 100% is the ideal case.

Figure 3a shows the calculated metrics U vs C. Centering value was often close to 100% using LSCM. However, uniformity values have a much broader distribution. Wide-field setups show a hig dispersion of C and U values. Finally, the analyzed spinning disk setups had less dispersed values closer to the ideal 100%. A closer look at this graph highlights interesting features, as indicated in Figure S3. For instance, most of the uniformity variations observed with confocal systems are associated with 405 nm excitation (Figure S4). Most systems designs involve the 405 laser line passing through a different fiber/lightpath than “visible” laser lines. The alignment is different and explains these differences. Moreover, both metrics associated with spinning disk systems show high values and lower dispersion (laser sources well aligned and centered). Finally, when using detector arrays, the sensor size influences the observed Uniformity values. Field illumination images acquired with large FOV (e.g. 13.312 mm × 13.312 mm) sCMOS cameras, often have a lower Uniformity (as most of the optics of the microscope were originally designed for the use of smaller imaging area CCD (e.g., 8.77 x 6.6-mm) or EM-CCD (e.g., 8.19 x 8.19 mm) devices. We indeed observed that low Uniformity/high Centering accuracy combinations are mostly associated with larger sensor sizes (magenta circle in Figure 3b). However, this is only the case for a subpopulation of sCMOS images as most of them show both high Uniformity/Centering accuracy values. Finally, the only “low Uniformity” and “low Centering” combinations (blue line of Figure 3b) were associated with misaligned TIRF microscopes, on which proper laser alignment has to be manually set before each experiment.

**Figure 3:**
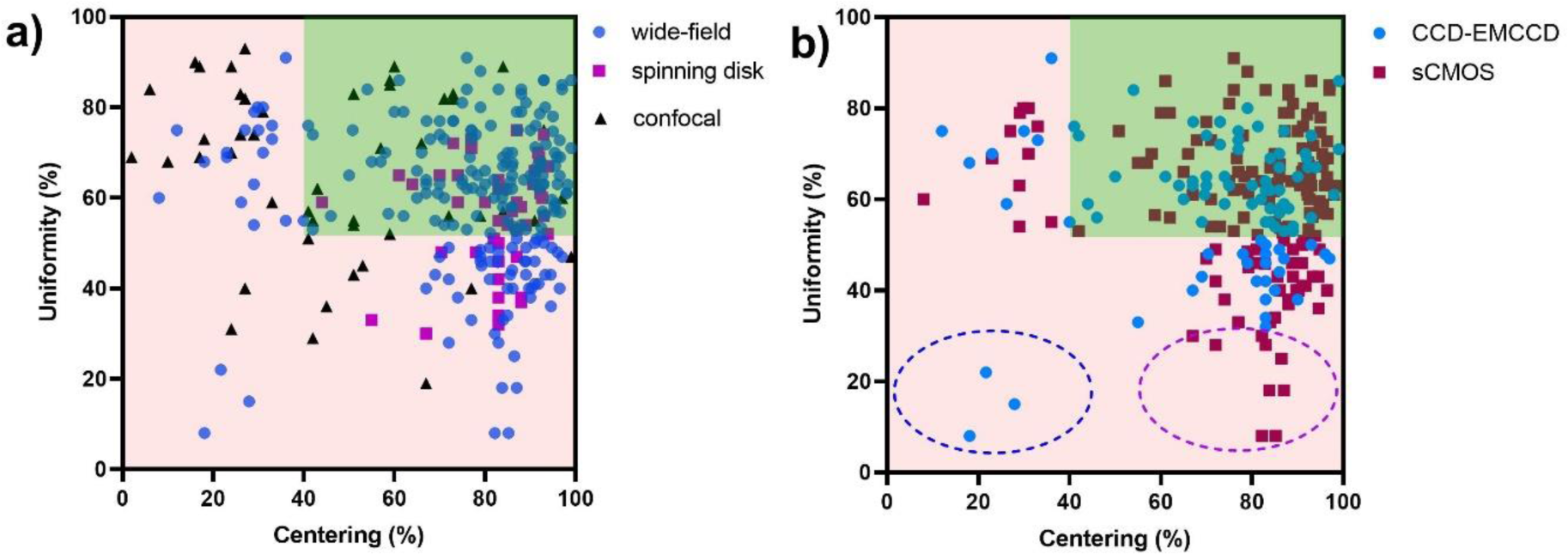
Distribution of Centering accuracy and Uniformity field illumination flatness metrics. Measurements were either classified using microscope type categories. (a) or two categories based on small (either typical CCD or classical EMCCD) or larger sensor sizes (sCMOS) (b). The blue dashed circle shows three values measured with TIRF microscope/EMCCD camera images. The magenta dashed circle highlights a subpopulation of low Uniformity/high centering accuracy couples, only associated with sCMOS camera images.

The third metric for characterizing the field uniformity uses the standard deviation of eight specific points of the image (i.e. the four corners and the middle of the four edges of the image). We further normalized this deviation using the maximum intensity found in the image and transformed this standard deviation to a 100% metric (100-NormalizedDeviation*100). The obtained metrics is called frame Uniformity (fU). The ideal case is when fU=100 (no intensity differences among the 8 points of the image). We found a similar fU dispersion compared to the Uniformity metric (Figure S5b). However, the uniformity metrics appear more sensitive (compared to the spinning-disk Uniformity/fUniformity spread in Figure S5a and 3a). We decide to stick to the use of both Uniformity and Centering accuracy values to characterize the field illumination quality.

#### Co-registration

Our co-registration study collected data from setups of the three microscopy techniques using over 70 different lenses. While sub-resolution beads are absolutely needed for resolution assessment, co-registration can be done with beads whose sizes range from sub-resolution (0.1 μm) to a few microns. We first studied the influence of the size of the beads on the co-registration results. We observed that beads with a size of 1 or 4 μm are ideal for co-registration studies, while co-registration results based on smaller 0.5 μm and 0.2 μm beads give a higher dispersion (data not shown). We believe it is harder to achieve a high SBR, as needed to further correctly identify beads and estimate their center coordinates, using these tiny beads.

We calculated the ratio of experimental to theoretical distance of the bead center coordinates for all two channel combinations (e.g., when analyzing the usual four channel images, six combinations need to be considered). Figure 4a shows the ratio distribution associated with blue-green (B-G) and green-red (G-R) channel combinations.

**Figure 4:**
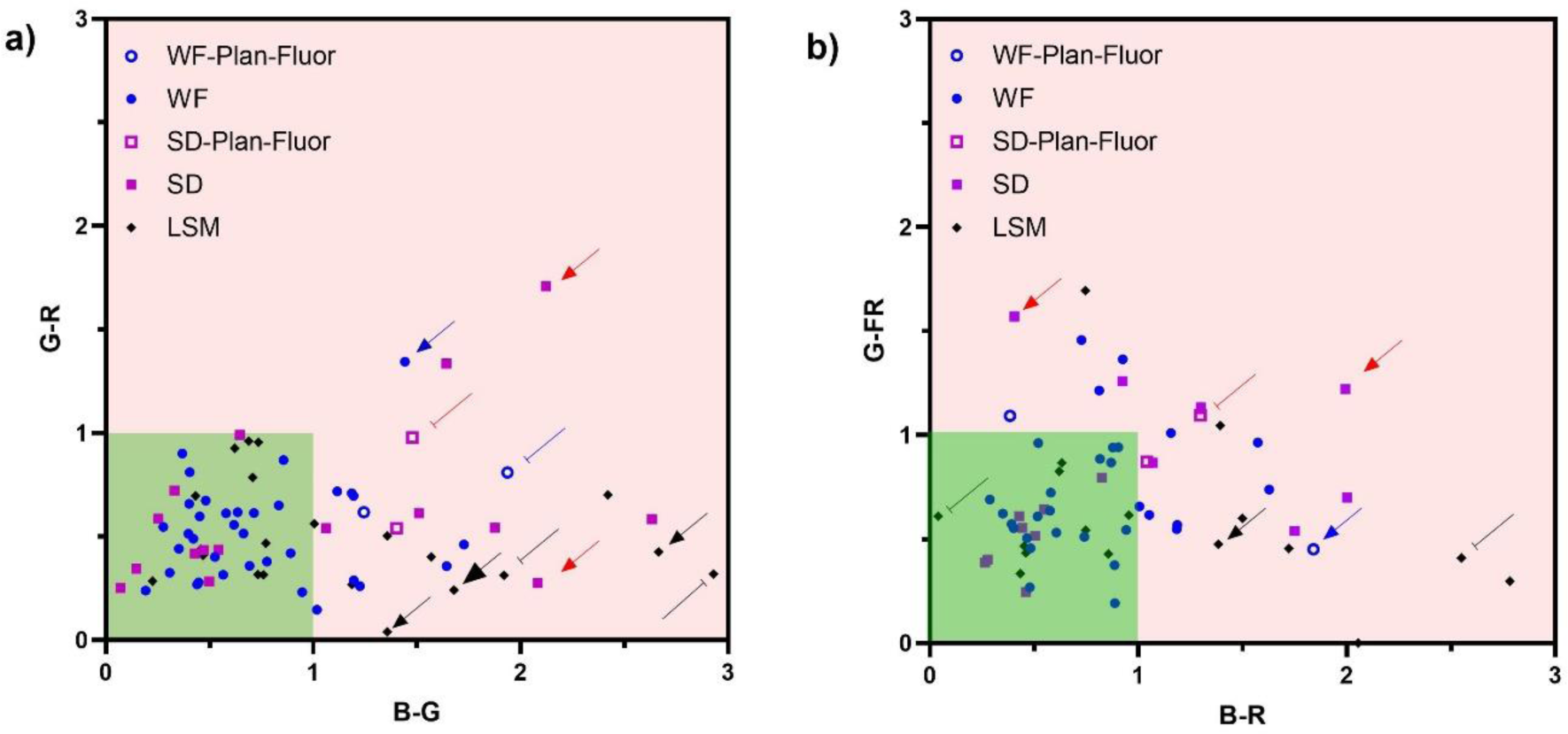
Co-registration distribution. a) Co-registration ratio of blue-green channels combination compared to green-red channels combination. b) Co-registration ratio of blue-red channels combination compared to green-far red channels combination. The green area highlights perfect co-registration in both combinations.

We observed that more than 75% of the results show nearly perfect co-registration in both channels, as both ratios are less than 1 (green zone in Figure 4a). When looking at individual combinations, 96% of the measures were below 1 for the G-R, 76% for G-FR, 67% for B-R, and only 63% for the B-G. This shows that most of the lenses used are better corrected for the G-R combinations, while the combinations with the Blue are more often associated with poor co-registration.

A comparison of the three different microscopy techniques shows that the above statement is true with all setups. However, spinning-disk confocal systems are more often associated with insufficient chromatic correction with 63% of the measured ratios below 1, compared to LSCM or wide-field workstations with 70% and 84%, respectively.

A closer look at outlier values highlights interesting features. The blue-caped line points to the round hollow blue spot in Figure 4a (e.g., B-G ratio: 1.94 and G-R ratio: 0.81) that was obtained with a 40x, 1.3 NA, Zeiss PLAN-Neofluar lens of a wide-field setup. This is in line with the incomplete chromatic correction of the achromat lens type compared to better corrected apochromatic objectives and some imperfect spherical aberration correction observed in blue PSFs (Figure 3). When looking at the spinning-disk configuration, we observed a similar difference. The red-caped line points to the square hollow red spot (B-G ratio: 1.48, G-R ratio: 0.98) that was obtained with a 40x 1.3 NA Plan-Fluor

Nikon objective. This objective is less corrected for chromatic aberrations between the DAPI channel and the rest than the Plan-Apo series and the B-G ratio is higher compared to apochromatic lenses. Finally, when using raster point scanning confocal, the black barrow (B-G ratio: 1.67 and G-R ratio: 0.24), associated with a 63x 1.4 NA Leica Plan-APO objective, shows that both objective combinations involving 405 nm excitation have a ratio higher than 1. Once again this issue may find an explanation with objective specificities. Hence, this objective is not recommended, and better-corrected lenses (lambda blue for Leica with better correction for the near UV light), should be preferred.

Observation of co-registration ratios can also be valuable to tackle other non-objective associated issues. The blue arrow (B-G ratio: 1.44 and G-R ratio: 1.34) was obtained with a 63x, 1.4 NA Zeiss objective Plan-APO of a wide-field system. After multiple tests, we found out that the filter set cube was not correctly aligned. A similar cube equivalent case was associated with 0.19 and 0.23 B-G and G-R ratios, confirming that the issue came from the misalignment of the cube components (dichroic mirror and filters) and was not related to some type of poor objective correction.

Another example is highlighted with the red arrows of 100x, 1.4 NA, and 40x, 1.3 NA objectives, respectively (spinning-disk). All objectives of this system have high values, far from the theory. Both objective chromatic performances were further investigated on a wide-field microscope to check whether the issue was associated with some objective imperfection or some misalignment within the light path. We found BG/GR ratios of 0.86/0.87 and 1.22/0.26 for the 100x and 63x lenses respectively. Hence, the system’s observed chromatic shift and poor chromatic performance are likely due to some other component misalignment upstream/downstream of the light path. It is sometimes easier to pinpoint the origin of poor chromatic performance. The black dashed arrows (B-G/G-R ratios of 2.93/0.32 for a 40x 1.3NA lens and 1.92/0.31 63x 1.4NA objectives) were taken with a Leica SP5 confocal, equipped with the classical light path design involving two laser injection fibers, one for the 405 nm diode laser and another for all other “visible” wavelengths. As correction mismatch was only observed with a combination involving the 405nm-associated B channel, it was easy to identify some 405nm light path correction lens, located before the objective, as the origin of these poor performances. The observed shifts were reduced after careful correction lens realignment (2.67-0.43 and 1.35-0.04 for 40x and 63x lenses respectively, black cap end lines).

Finally, Figure 4b shows the co-registration results for two different color couples (blue red B-R and green far-red G-FR). Compared to Figure 4a, we observed a similar dispersion for the B-R ratio metric of the couple that presents many values higher than one. The G-FR couple is slightly worse than the equivalent G-R that is expected as the wavelength difference is higher. Using the same arrow type/color scheme, most of the extreme cases highlighted in figure 4a are outliers in figure 4b, and their values are mostly far from 1 as in figure 4a.

#### Illumination power stability

Different time scales for monitoring the illumination power stability can be used. For short-term and mid-term monitoring, fluctuations were recorded during a single experiment, either 5 minutes or 2h long, respectively. These settings are typical for 3/4D stack acquisitions of either a fixed or live sample. Monitoring over months-long periods helps to characterize the light source intensity properties during the lifetime of a microscope.

Before defining the metrics, a handy first approach is to characterize the measurements qualitatively. Figure 5a and Figure 5b show the mid-term monitoring of two different raster single point scanning confocal microscopes after a 1h long warming-up delay. Both microscopes have either gas lasers (488 nm Argon and 633 nm He-Ne) or diode/solid-state lasers (diode 405 nm and DPSS 561 nm). To define the appropriate warming-up time, that will further be important for accurate intensity quantification, power was measured for 3h starting directly after the laser was switched on (Figure S6). Diode-pumped solid-state lasers proved relatively stable within a short time, while gas lasers or 405nm diodes needed some more time to stabilize. This unexpected longer 405 nm diode warm up period was quite a surprise and might be explained by some other optical/electronic component within the 405nm beam path (as power was observed either at the back focal plane or at the focal plane of a dry 10x lens). Figure 5b (Leica SP5 setup) shows i) some decrease of the DPSS 561 nm laser power (as compared to a stable Zeiss LSM880 setup) and ii) high fluctuation of the power of the Argon 488 nm line. These observations were a warning for the facility team, who after contacting the service technician of the microscope manufacturer and after a visit on site, managed to identify and correct the problem. A damaged fiber and aging of the lasers were identified as the main reasons for these poor performances.

**Figure 5:**
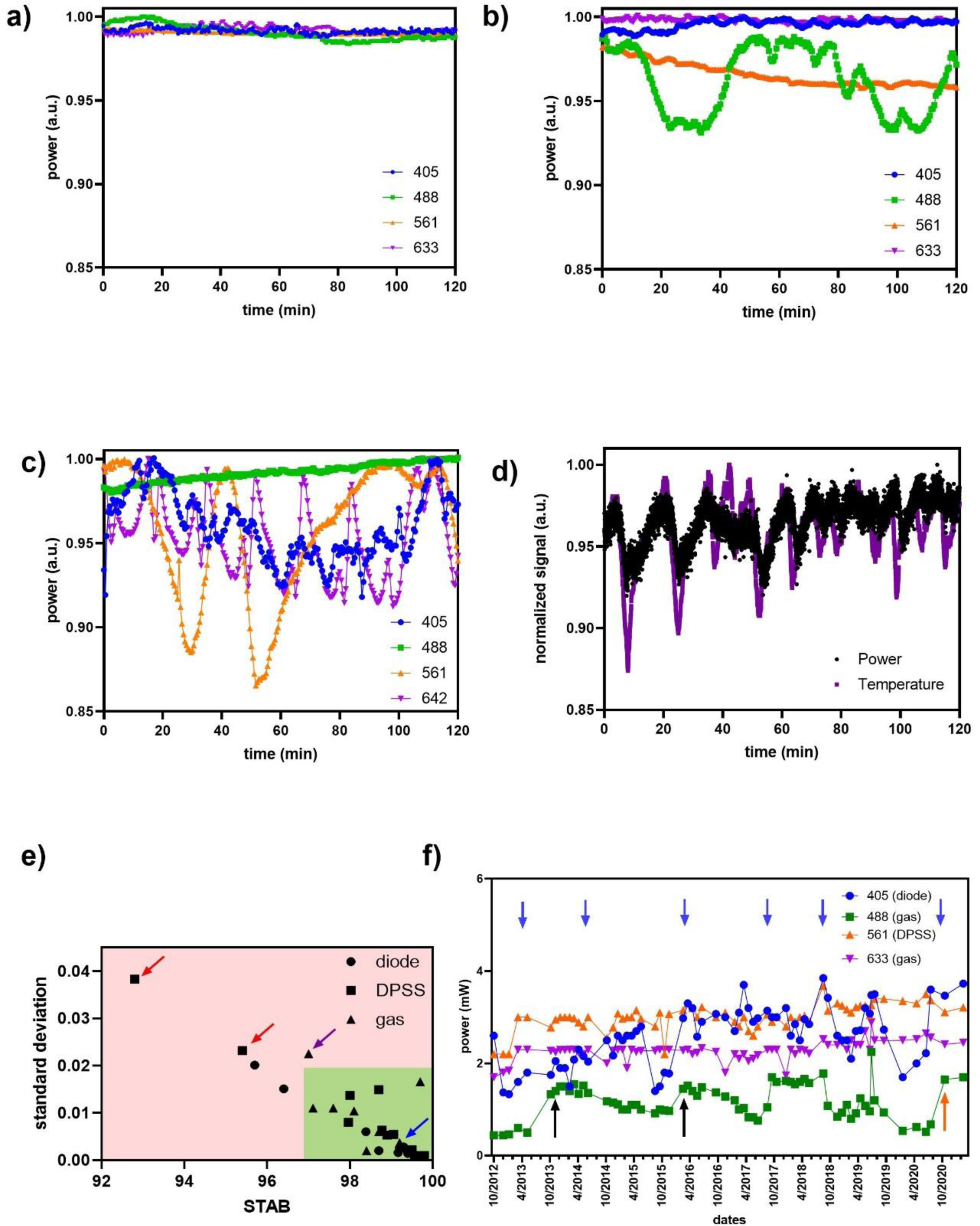
Illumination power stability. a) Power stability over 2h for a confocal LSM 880. b) Power stability over 2h for a confocal SP5. For a, b the 488 and 633 lines are gas lasers. c) Power stability over 2h of a TIRF microscope. d) Power of 561 nm laser of (c) and room temperature stability over 2h. The minimum and maximum value of temperature was 19.5 and 22°C respectively. e) Laser power mid-term stability STAB vs. standard deviation of normalized to 1 (max value) laser power values over 2h. The blue arrow points to the 488 laser of (a), the magenta arrow points to the 488 laser of (b), the red arrows points to the 561 and 642 lasers of (c). f) Illumination power monitoring over a microscope SP5 lifetime (long-term stability). Intensities of a diode 405nm, 488nm Argon & 633nm He-Ne gas lasers and a 561nm DPSS laser were recorded during nine years. The Argon 488nm laser was replaced twice (black arrows). The orange arrow indicates when the optical fiber of the 488 nm laser was replaced. The manufacturer services are indicated with blue arrows (and are often associated with power increases as the system is realigned).

Some other aberrant behavior can also be observed, as indicated in Figure 5c (mid-term monitoring of a four solid-state laser TIRF microscope). While the 488nm laser is extremely stable, all other wavelengths showed erratic cycle-like fluctuations. The underlying reason for that was an irregular polarization shift in the optical fiber, influenced by temperature instability of the room. To ensure the temperature instability hypothesis, we performed simultaneous power and temperature measurements over 2h, every second this time. The temperature was measured at the optical fiber level. Figure 5d shows that there is a correlation between the power and temperature fluctuations.

To quantify the laser power fluctuations, we use two metrics. The first metric is the standard deviation of the power measurements, which are normalized to 1 for all monitored lasers. The second metric is the STAB factor, which considers the minimum and maximum value of the laser power and is defined in the ISO 21073:2019 norm^13^. Figure 5e shows the metrics values for the measured laser stabilities. The lasers were classified into three categories: diode lasers (including OPLS, i.e., the lasers that can be controlled directly and do not require any modulating device such as AOTF or AOM), DPSS lasers, and gas lasers (e.g., Argon or He-Ne lasers). Most lasers have high STAB values and low standard deviation metrics (Figure 5e, green zone), including the stable Argon 488nm laser line of Figure 5a. Some other lasers have lower STAB (<97%) and higher standard deviation (0.2), such as the Argon 488nm laser line of Figure 8b. Extreme cases (e.g., 561 nm laser of Figure 5c, red arrow of figure 5e) show very low stab and high fluctuations/standard deviation.

A short-term analysis of a subset of these lasers gave similar results (Figure S7). As the behavior of the lasers of the same setups (blue and closed orange arrows) are similar, this suggests that short time scale stability issues are likely linked to the stability of common electronic and/or optical components that all lasers light finds until reaching the objective plane, rather than laser-related individual issues.

A third long-term, monthly monitoring of the light power indicates fluctuations that can be observed and should help to compare different experiments acquired at different periods. A decrease of the specific label emissions can be due to laser misalignment and sudden power decrease and, hence, would be unrelated to the sample itself. The core facility staff or the microscope user should thus regularly monitor and record the laser power. Figure 5e shows the long term stability of four laser lines on a scanning point Leica SP5 confocal over nine years. This typical system was fully covered by the manufacturer aftersales service, and, as a consequence, access to any alignment tool was restricted to manufacturer staff. As expected, technician visits or whole laser replacements had a positive effect on illumination intensities. As highlighted in Figure 5e, lasers are more or less stable, and long-term monitoring proved necessary to achieve the best system performances/stability.

#### Stage drift and positioning repeatability

##### i Stage drift

The stage drift characterization is a fairly straightforward procedure to follow. However, sources of drift can be multiple and difficult to identify. The assessment involves some self-made bead slide with fixed stable beads. A fluorescent bead is convenient, but brightfield observation of various features is possible as well. 3D time-lapses were acquired every 10 min over 15h. If present, hardware and software focus correction(s) should be deactivated to allow raw 4D drift evaluation. Individual beads were identified and tracked over time. 3D bead coordinates and the drift speed along each axis were calculated. A maximum 4D projection of a typical drift experiment, over 90 time points, and the corresponding bead track are displayed in Figure 6. The 3D displacement decreases over time (Figure 6b), suggesting a stabilization. Figure 6c shows the calculated displacement along each axis. An initial strong y-axis drift was observed, whose amplitude decreased after some time. Z-axis drift was negligible, for standard microscope experiments. Bead maximum intensity projection allows a first qualitative approach, as it shows a bead drift along the y axis (Figure 6a). When considering the maximum acceptable drift, a timeframe and a minimal displacement should both be defined. Here, we considered any axis displacement within 10 minutes should be kept below a high NA minimal resolution distance of 200 nm. That drift should be barely noticed during 10 minutes in diffraction-limited conditions. This means a stage would be considered not affected by drift if all three-axis displacement speeds are less than 15 nm/min for each axis. The stabilization time in Figure 6 was 105 minutes. Before stabilization, the mean 3D speed of the system is 56 nm/min, and after stabilization, 8 nm/min. One should consider that this stabilization time depends mainly on environmental conditions, particularly related to temperature variations that can imply large drifts.

**Figure 6:**
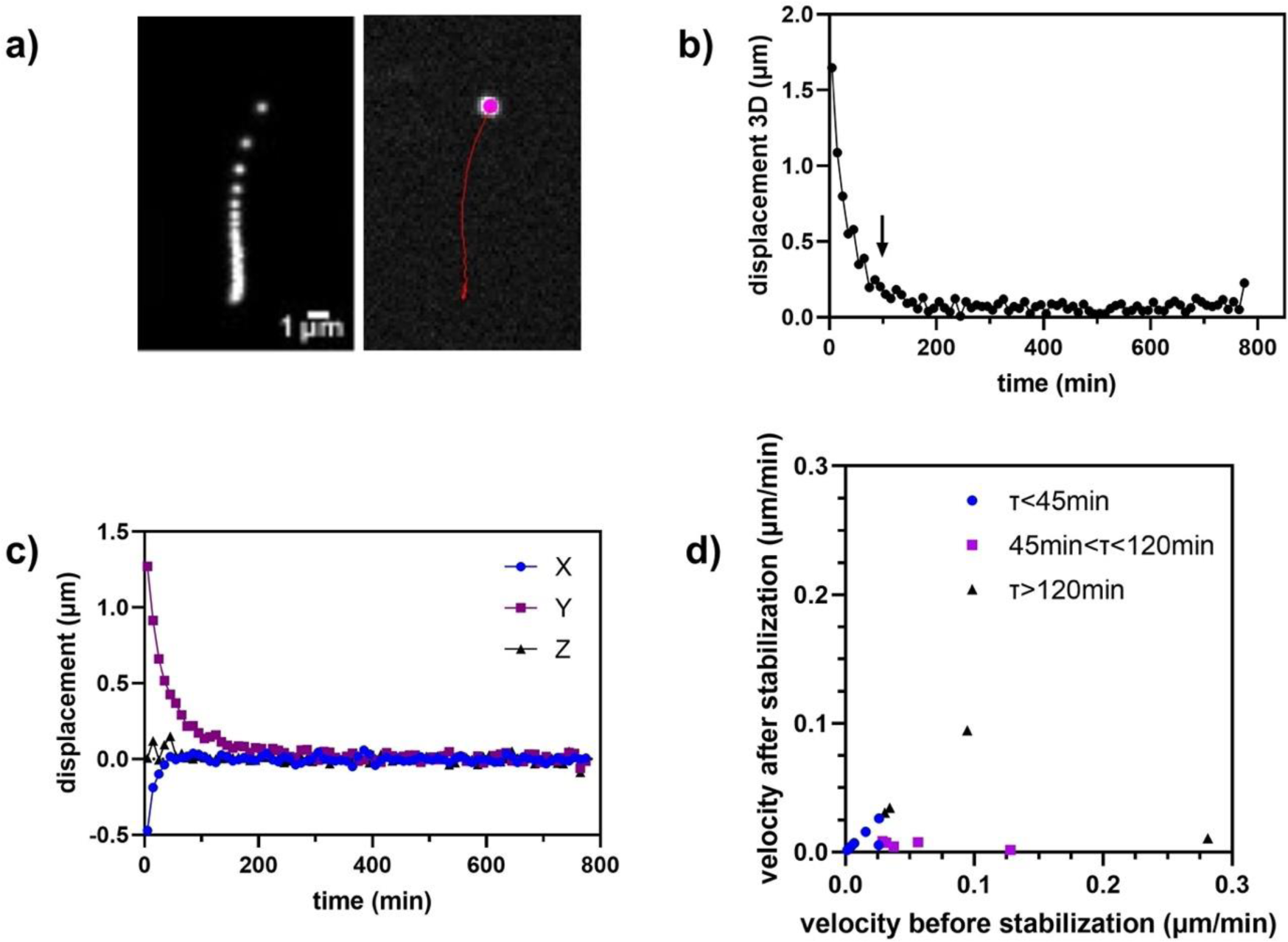
Drift measurement. a) max Z projection of the time projection of a time-lapse for 15h of a fluorescent bead, and trajectory of the bead, with the initial segmented bead position shown in magenta, b) Global 3D displacement over time. The arrow indicates the calculated stabilization time. c) Relative displacement along the three axes. d) calculated velocity before and after stabilization of different microscope stages grouped in 3 categories upon their stabilization time τ.

Figure 6d displays a distribution plot of 3D drift speed before and after the calculated stabilization times of 16 microscopes. When short stabilization times (<45min) are observed, both rates are almost identical. However, for longer stabilization time (45min< τ < 120min), as well as for very long stabilization time (>120 min) the two speed values are different. Finally, sometimes no stabilization was found during the whole 10h time-lapse.

We then investigated the drift speed fluctuations over days. Three different measurements on the same system were performed during three consecutive days. The drift values were very close, and the velocity dispersion low (data not shown). However, during a month or even a year, drift of a system can change significantly if environmental parameters change, like the room temperature or even the temperature flow in the room. Time monitoring of the drift is then necessary if such changes are frequent.

Using the three metrics and this experimental dataset, drift behavior can be classified into three categories according to the specificity experiments and its need for precision (Table 1). When the velocity before and after stabilization is identical and really small (lower than 15 nm per minute, which is the theoretical resolution of a high NA objective lens after 10+ min), the drift can be considered negligible (high precision experiments, e.g. super-resolution purposes). When both a 45- 120 minutes stabilization time and 3D velocities within a 15-50 nm per minute range are observed, drift can be considered acceptable (e.g., time-lapses at the cells scale). Any other drift parameter combination should be rejected or taken into account during the image analysis step because it can lead to serious misinterpretation.

**Table 1:** Drift tolerances. When the velocities before and after stabilization are identical and kept low, the drift is negligible. When slight velocity differences are observed while the stabilization time stays short (τ<120min) the drift is acceptable. When there is a big velocity difference associated to a high stabilization time or any other case (such as no stabilization), the drift should be considered unacceptable.

| Drift | Unacceptable | Acceptable | Negligible |
| --- | --- | --- | --- |
| Mean velocity after stabilization | >0.05 $\mu\text{m}/\text{min}$ | [0.015;0.05] $\mu\text{m}/\text{min}$ | <0.015 $\mu\text{m}/\text{min}$ |
| Mean velocity before stabilization | >0.1 $\mu\text{m}/\text{min}$ | [0.015;0.1] $\mu\text{m}/\text{min}$ | <0.015 $\mu\text{m}/\text{min}$ |
| $\tau$ : stabilisation time | >120 min | [45;120] $\mu\text{m}/\text{min}$ | <45 min |

##### ii Stage positioning repeatability

Microscope stage specifications include maximum travel range, (maximum) speed, repeatability, or accuracy. When acquiring multiposition experiments, a key parameter is the stage repeatability, i.e., the system’s ability to reposition to the same point. The ideal stage should have high accuracy and repeatability. The two parameters are closely related but independent. Systematic errors that influence stage accuracy can be taken into account and compensated for, but after such corrections, repeatability will be the ultimate limit of the device.

The repeatability is calculated as the standard deviation of the position coordinates during successive moves to a set of predefined positions. It can be unidirectional (the stage returns to a given point with only one axis movement along x or y) or bidirectional (the stage returns to a given point coming from a random previous point, involving x and y movements).

Repeatability can be improved when a backlash correction is applied and/or when the stage has linear encoders. The backlash is the systematic error created by lost motion in the drive mechanism that appears when changing direction. An anti-backlash algorithm is often used to correct this effect. When x-y linear encoders are mounted on the x-y plates of the stage, then the repeatability is further improved.

Figure 7a shows the repeatability values for the x-y axis for 16 different stages on 16 microscopes. Stage suppliers were either Märzhäuser or ASI, and the majority of the stages had linear 2 phase stepper motors. Two of the stages had linear encoders (marked with an arrow or a caped line).

**Figure 7:**
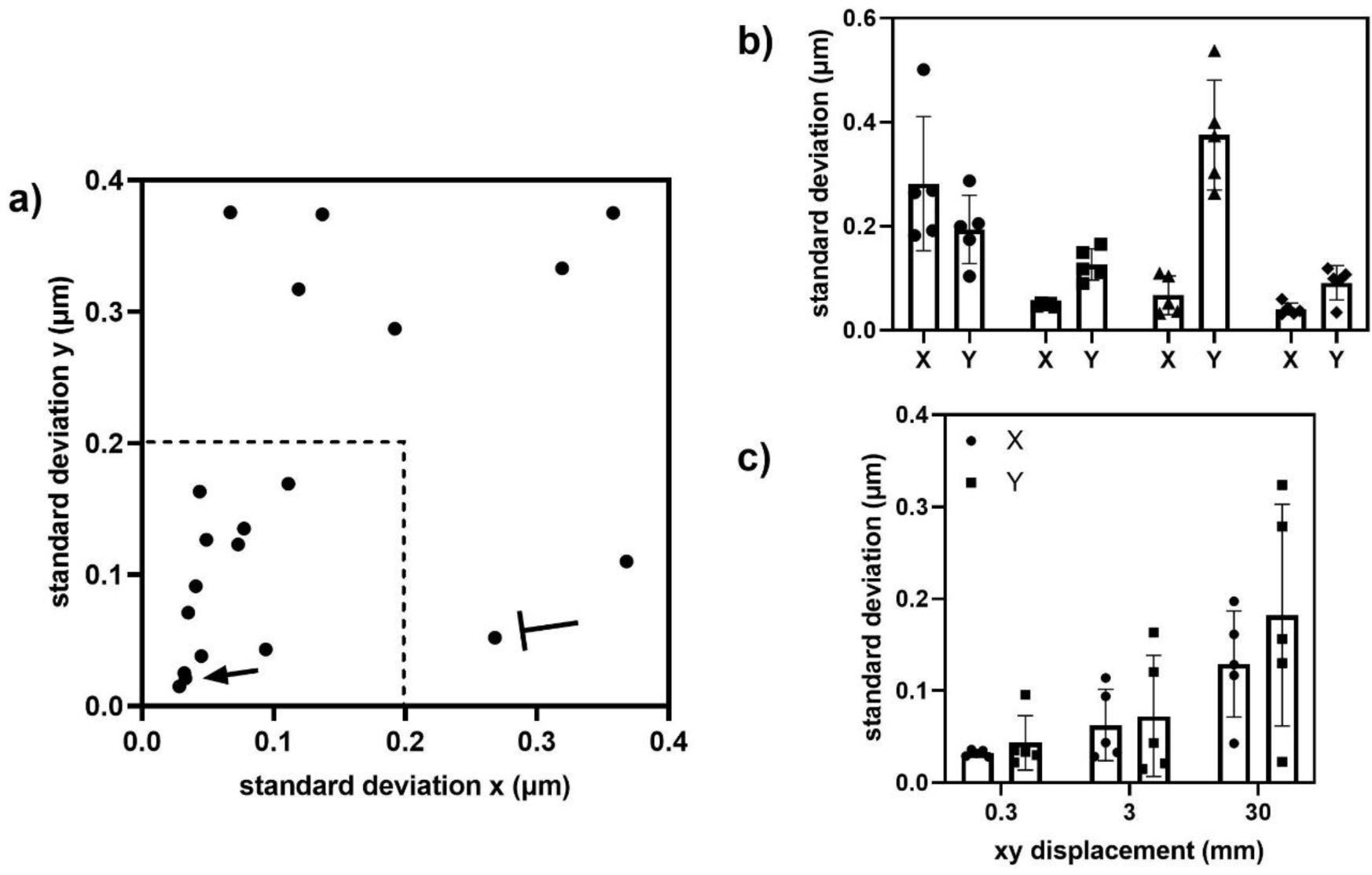
Repeatability characterization. a) Bi-directional stage repeatability for 3 mm in both axes (4.2 mm diagonal) stage displacements. The arrow and capped line show two linear with encoders same model stages mounted on the same microscope. b) Reproducibility in time of stage xy repeatability measurements for four stages. c) Influence of the stage displacement on the repeatability.

Measurements during 20 time points were performed from the center to a distance of 3 mm at x and y and back at the four corners of a square. The variability σ of the measurements of the central point gives the repeatability values along the x and y-axis. The variability was always below 0.4 μm, within specifications (i.e., below the usual maximal 1 μm manufacturer’s specifications). A distribution analysis highlights a population with values below 0.2 μm at both axes (zone in dashed line in Figure 7a). It is not easy to define limit values for repositioning, as it highly depends on the stage type. For instance, for the two ASI stages having linear encoders, although they have the same characteristics and were mounted on the same microscope, they present different repositioning values. After the repositioning measurements, we identified that the first stage was out of the specifications (caped line of Figure 7a) and it was replaced by a second one found in the specifications (arrow).

We examined some parameters that could influence repeatability. The stage speed did not significantly affect the values (data not shown, varied rates from 1 – 22 mm/s). We examined the reproducibility of the measurements by performing acquisitions for five consecutive times. We observed that there was almost no variability in some cases, and in other cases, it varied a factor of two. Figure 7b shows the results for four different stages. This is a strong indicator that the repositioning measurements do not depend exclusively on the stage characteristics but the entire system stability.

We examined the influence of the stage displacement distance using three different values (0.3 mm, 3 mm and 30 mm) for both axes for five different stages from the same constructor (Märzhäuser) and with similar datasheets. A 3 mm distance is often a displacement used for multi-position experiments within the same well/coverslip. Longer lengths (e.g., 30 mm) are typically used for multi-well samples. We represent in Figure 7c the mean values with their standard deviations for both x and y calculated standard deviations for the five stages. Although there is a high variability (measurements not made on the same microscope and the stages were not exactly the same model) we can distinguish that the repeatability gets worse for longer displacements.

Other parameters can influence the repeatability here, like the proper fixing of the sample, the waiting time (if it exists) for each position, the acceleration of the stage and the temperature variations.

##### Camera noise characterization

The read noise was calculated from 3 CCD, 4 EMCCD, and 22 sCMOS cameras from various constructors and most often for both standard and fast camera readout modes (30, 100, 200 or 500MHz for sCMOS, 10 and 17 or 30MHz for EMCCD, 10 and 20MHz for CCD). Read noise maps and read noise distributions are indicated in Figure 8a, b. The difference in the measured read noise and DSNU value is shown in Figure 8b when the blemish correction (warm pixels or defects) is ON or OFF. Extreme pixel values are smoothed with the nearest neighbors, and the calculated noise is closer to the one in the datasheet. For each sCMOS camera, the dark offset and the DSNU distribution map were also calculated (Figure 8c). Data for each pixel was averaged for 100 frames, and a narrow look-up-table was used for visualization to render small differences in the columns visible.

**Figure 8:**
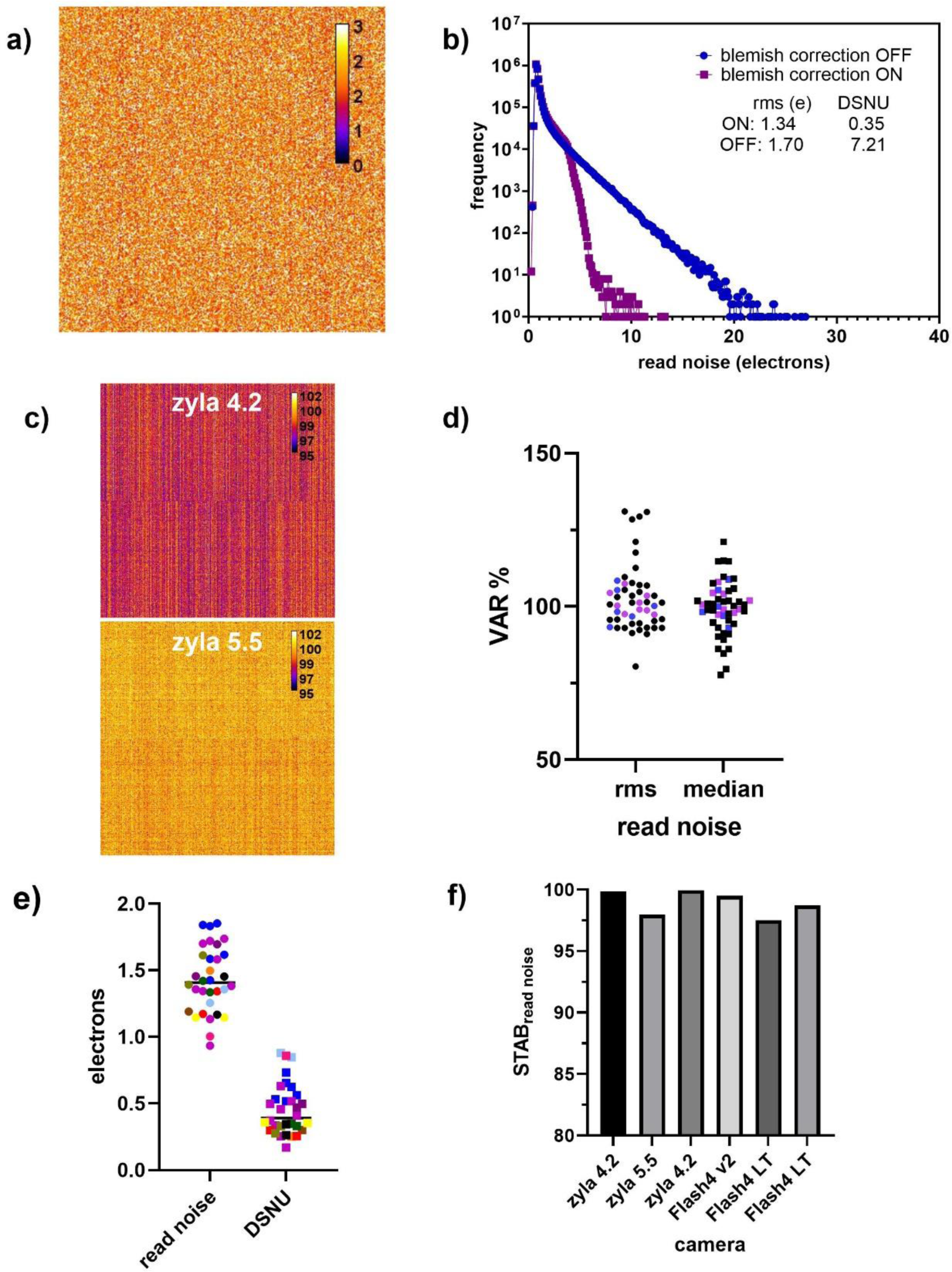
Read noise and DSNU characterization of cameras. a) read noise map of an Orca Flash4 v3 (zoom of 300x300pixels). Scale bar in electrons. b) Comparison of blemish correction on read noise for the camera of a). Read noise distribution for pixel correction ON (blue) or OFF (magenta). Inset table: measured values of read noise (rms) and DSNU for both cases. c) DSNU map (dark offset of 100 images averaged) for two different cameras in the central part (500x500 pixels). Different patterns are distinguished, depending on the sensor corrections (column, pixel) and the vertical juxtaposition of the two matrix of this kind of sCMOS. Scale bar in digital number (DN). d) Read noise rms/median VAR metric distribution for CCD (blue), EMCCD (magenta) and sCMOS cameras (black). e) Read noise (rms) and DSNU distribution (as expressed in electrons) of sCMOS cameras (each colour corresponds to a single camera, with different readout speed modes). f) Stability metric (STAB) of the read noise for six cameras over six months.

Figure 8d shows the VAR metric distribution for both rms and median read noise values. We calculate this metric for both noise values as camera constructors often provide a single readout noise value (either rms or median), mainly for commercial reasons. The majority of the VAR values are close to 100%; 84% percent of the calculated rms values are within 10% variation from the 100% value. This shows that measured readout noise values are close to the manufacturer’s specification. Some of the values are higher than 100% (better noise evaluation compared to specifications) and were most often observed when a blemish correction was applied. For CCD and EMCCD cameras, the rms or median readout noise values are almost identical and VAR distribution is kept in a narrow 10% variation window.

Figure 8e shows the rms read noise and DSNU of the 22 sCMOS cameras in electrons included in the Figure 8d analysis. The calculated DSNU is never higher than the read noise, as expected from theory for this kind of cameras.

For precise DSNU evaluation, one should in principle acquire thousands of frames to eliminate the read noise. To define the optimal dataset size, we introduced a quality criterion that compares the DSNU obtained with datasets of different size. A threshold of 10% yielded an optimal dataset size of 100 frames (i.e., the DSNU value difference obtained with larger datasets is less than 10% of the DSNU value calculated with 100 frames, Figure S10). Hence, we recommend using a 100 dark frames dataset for DSNU evaluation.

The evolution over a six months of read noise values of different cameras, as measured with the STAB_noise_ metrics, is shown in Figure 8f. STAB_noise_ was always higher than 97%, meaning that camera read noise is relatively stable, and any fluctuation indicates serious detector issues.

## Discussion

### Microscope Lateral and Axial Resolution (PSF)

The analyzed dataset was collected across ten microscopy facilities, 91 objective lenses of various magnifications and corrections and a wide range of instruments. The PSF measurements were at an average of 1.54x, 1.52x and 1.31x larger than the lateral, theoretical resolution for wide-field, single point scanning and spinning-disk confocal microscopes, respectively. At the same time, axial performances were 1.16x, 1.15x and 1.41x larger than the respective theoretical values (Figure 2b). The observed wide-field higher dispersion of the measured/theoretical axial resolution ratio, as compared to both confocal types, could be associated with some spherical aberration issues. Hence, the confocal effect at 1 A.U. crops the elongated PSF of a spherical aberration-affected objective, reducing the ratio distribution dispersion that such PSF induces in the wide-field mode. Voxel size limits accurate FWHM measurements. As proper, Shannon-Nyquist criterion compliant sampling rate is easy to achieve, PSF evaluation of an LSCM is not such an issue. However, wide-field and, even more, spinning disk microscopes involve a fixed camera pixel size, which can sometimes be quite high (e.g., some EMCCD), and the following under-sampling, as observed with 60/63x or 40x objectives, limits the FWHM estimation accuracy. For spinning disk microscopy in particular, the most often fixed pinhole size can also effect the measured deviation of the measured FWHM compared to the theoretical values. It should be noted that the theoretical values consider subresolution point sources. In our case we use 0.175 nm beads that give slightly larger PSF values compared to point source PSF (one should convolve the two PSFs for the theoretical values).

The resolution performance measurement is highly impacted by the objective quality. Furthermore, it strongly depends on additional parameters, ranging from the sample quality to the signal strength and the image noise. Variability is also induced by the user (whose capacity to fully follow a strict procedure may vary). Besides any lens-induced aberration, special care should be given to following the same acquisition protocols, using the same slides, and paying special attention to avoid errors such as forgetting a DIC prism within the optical path, not cleaning the objectives before the measurements, or not properly setting the correction collar, when necessary. We tried to limit the “user” parameter since only experienced facility staff carried out all measurements. However, although user-associated variability was kept low, one cannot entirely avoid it. Variability was most often associated with suboptimal sampling rate (e.g., inevitable for spinning-disks) or blue-emission dyes (e.g. DAPI). As the objectives and the whole optical path are usually better corrected in high-quality systems, adding aberration correction within the UV or near-UV wavelength range is challenging. The microscope user should use all available tools for troubleshooting. An adjustable pinhole at the LSCM offers such solutions. First, one has to be sure that the pinhole is well aligned before performing any confocal PSF measurements. Whenever the performance significantly diverges from theoretical values, we recommend using a fully-opened pinhole, or checking the objective on a wide-field system. These comparisons are helpful to precisely identify the origin of the poor performance (i.e., objective-associated or related to some other confocal component).

The method described by Zucker et al.^27^ using a metal-coated mirror and detecting the reflected light of the laser can be performed to ensure the pinhole alignment and measure the axial resolution in an alternative way.

We propose experimentally defined limit values, as extracted from the PSF FWHM distribution of Figure 2a. Objective resolution performances can be considered within limits if the lateral and axial measured/theoretical ratios are kept below 1.5. Any ratio above these values should trigger some in-depth investigation to point the origin of such poor performances. Although such limit values may seem quite high, one has to keep in mind that these tolerance values rely on some theoretical formulas, considering optimal conditions such as shot-noise free confocal imaging.

Comparison of x and y FWHM helps to extract valuable information on the PSF symmetry. While asymmetry was negligible in wide-field and spinning-disk confocal setups, the average asymmetry ratio of point scanning confocal microscope is 0.86. This is expected, as FWHM is smaller in the direction perpendicular to the direction of the linear polarization of the excitation laser in high NA objectives (NA>1.3) ^44, 45^.

### Field Illumination

We characterized the field illumination uniformity of various setups, using 145 objective lenses of different magnifications and various excitation/emission wavelength combinations. We recommend using fluorescent plastic slides, as they do not need special preparation. However, precise, quantitative characterization of the uniformity, as required for any shading correction for instance, should be performed using a slide loaded with a thin layer of a fluorescent dye^46^. Although we defined three metrics (Uniformity, Centering and flat Uniformity), since flat Uniformity was most of the time giving the same information as Uniformity, we recommend using only the first two metrics U and C.

For the Uniformity metric, the size of the wide-field detector chip has a strong influence, as often observed using large sCMOS sensors cameras (Figure 3b). Whereas smaller sensors have been used in the past, most manufacturers have recently introduced wider field corrections for objectives, and the whole internal microscope stand light path. Hence, whenever affected, besides choosing better corrected objectives, no corrective action can be undertaken for the microscope stand, apart from cropping the image to the central area of the sensor or using shading correction.

Defining experimental limit values for the field illumination is not trivial. However, we propose, as defined using experimental data from Figure 3a distributions, that Centering above 40% and Uniformity above 50% can be considered acceptable. As mentioned above, and in Figure S4, some corrective action may be possible or not. Moreover, both low Centering and high Uniformity values will be a warning, but the images stay exploitable somehow. However, low Uniformity values associated with any Centering value are not exploitable. Some action should be taken if the images are to be quantified (e.g., lasers realignment, cropping the sensor area, etc.). We conclude that both metrics are interconnected and, to have a uniform and centered field illumination, their respective tolerance values should be both taken into account.

### Co-registration

We studied the co-registration of 80 objectives, across the three microscopy techniques. 74% of the measurements yielded co-registration ratios below 1 (meaning the measured chromatic shift is below the resolution limit). G-R combination was the best-corrected (96%) compared to G-FR, B-R, and B-G (75%, 67% and 62% respectively). This clearly shows that, when observed, shifts are likely to be associated with imaging of near UV excited dyes. Whenever these are part of the channel combination, ratios are below one in 78%, 47% and 58% using wide-field, spinning-disk and LSCM setups, respectively. 85% of the ratios of the other pairs were below 1 (such as G-R and G-FR). Wide- field microscopes performed apparently better as compared to confocal (90%, 85% and 78% for wide-field, LSM and spinning-disk respectively), as observed with combinations involving near-UV.

This may come from a lower theoretical resolution for the wide-field microscope, making the ratio slightly lower. It may also come from the lasers needed by LSCM and spinning-disk microscopes that have to be carefully aligned and checked to ensure proper stability. In contrast, an alignment of a wide-field microscope is much simpler. For instance, the misalignment of the UV lens in the confocal showed extreme values and triggered a very high ratio for near-UV images. The realignment of the UV lens was very efficient for the 63x objective but not for the 40x, meaning that one intermediate position has to be chosen or preferred or has to be adapted each time for the specific objective. For wide-field microscopy, the positioning of the dichroics in the filter cubes can influence the channels’ co-registration. Spinning-disk microscopes were the most affected during our studies, suggesting that their design was more subject to chromatic shift. It might involve that the different wavelengths have more chromatic shift or are less corrected while going through the spinning-disk unit and the microlens array of the disks. In any case, the most adapted wavelengths for live-cell imaging (green – red emission channel) were quite well aligned, showing that the microscopes with all their components are most adapted for imaging at that range of the visible spectrum.

### Illumination power stability

Monitoring illumination intensity is a key element to ensure that the comparison of fluorescence intensities makes sense. While illumination intensity can be measured at any point of the light path, including directly after the light source, it is advisable to monitor it at the objective focal plan such that all parameters influencing illumination intensity are integrated, and the actual luminous flux received by the sample gets measured. Short and mid-term fluctuations, i.e., within the time frame of a single experiment, can bias quantification. While comparisons over more extended periods are quite challenging, as much more parameters may also affect the experiment (e.g., sample changes, sample preparation variations/modifications). Monitoring such fluctuations is indicative of the “health” of the setup. Knowing these long-scale fluctuations also gives hints on how to correct data to achieve some qualitative, rough comparison. Using two metrics, the standard deviation of the normalized intensity and the STAB (stability) factor, we show that once the system is warmed up, any further fluctuations are likely to arise because of issues such as laser aging (especially for gas lasers), inadequate polarization stability, or misalignment of elements such as optical fibers. Defining the light source warming time is not trivial, as some lasers show a warming period of some hours (Figure S6). Based on our experimental data, we proposed a tolerance value for STAB of 97%, below which the source is considered as unstable, and a normalized intensity standard deviation limit of 0.2, above which fluctuations are too high. Whenever measured values are out of these limits, then careful identification of the origin of those fluctuations should be undertaken followed by appropriate corrective actions.

We focused our studies on lasers. Most measurements involving solid-state lasers were within limits as soon as the stabilization time was reached. When above tolerances, other instability creating sources were found (e.g., insufficient fiber coupling, temperature variations). Gas lasers, typically aging Argon laser tubes, showed strong intensity fluctuations.

Fluctuations affecting source stability during the long-term monitoring of the illumination intensity involved aging of the source, optics alignment stability, fiber coupling, or some damage of optical elements (e.g., deterioration of an AOTF by a 405 nm laser line). Hence, whenever fluctuations are observed, we recommend, if accessible, a supplementary measurement straight at the laser output to check the stability of the source before going through all the optical elements. However, this is only possible in some cases and not compatible with a closed commercial system. Long-term monitoring is instrumental in deciding if a laser needs to be replaced. The above tolerance values may be tuned might need to be adjusted depending of the used laser type. Gas lasers tend to show high mid-time scale fluctuations and their aging is often associated with a global intensity decrease (Figure 5b shows the fluctuations). A drop by a factor of three is a good indication that the laser needs to be replaced or the tube current to be adjusted. However, a third of the original value is usually still sufficient for most experiments (excluding FRAP). An observed illumination intensity decrease associated with a solid-state laser is likely to be linked to some misalignments.

### Stage drift and positioning repeatability

#### i Stage drift

The stage drift measurements are quite easy to perform but their interpretation can be more complex. The measured drift depends on various parameters such as mechanical and temperature stability. For instance, the same microscope with the same stage can give completely different drift values when placed in another room, with different temperatures, humidity, or air flow variations. We applied a protocol to measure the drift during overnight experiments easily. The temperature at the sample level was considered stable. Different metrics were calculated, including the stabilization time and the 3D drift velocity before and after stabilization.

Three categories were defined according to combinations of stabilization time and velocities values. Drift may be considered negligible whenever the measured stabilization time is <45min and both before and after stabilization velocities are kept low (as defined by a subresolution 200 nm drift during 10 minutes). When stabilization occurs within 45 and 120 minutes and the 3D drift is below 50nm/min after stabilization, the drift is considered acceptable, although it may affect image analysis and imply further post-processing. We then considered the drift as unacceptable if the stabilization time is >2hours and 3D displacement after this stabilization is higher than 50 nm/min, such values should trigger some in-depth investigation up to the stage replacement.

One has to keep in mind that drift measurements only make sense if environmental conditions (e.g., temperature, humidity, air flow, warming time of the stage) are kept stable. If these change (e.g., depending on the period of the year), the drift measurements are no longer valid. Hence, regular drift characterization is necessary when environmental conditions frequently change. A stabilization time is always required before starting the acquisition. Furthermore, it is very important to consider that once the sample is fixed on the stage, it is an integral part of this device. To preserve intrinsic heat of the stage, it is possible to leave the controller under permanent voltage, avoiding additional drift due to periodic stage heating cycles.

#### ii Positioning repeatability

Stage repeatability is directly linked to stage drift. If parameters involved in drift are controlled or characterized, one can measure the ability of the stage to reposition at the same X, Y location (repeatability). As drift has to be kept negligible, monitoring repeatability must be performed after the stage stabilization time. The sample should also be well fixed to avoid a sample-associated source of drift. Finally, experiments were performed with dry lenses to prevent any additional drifts associated with surface tension effects of the objective immersion, especially for long displacements.

Manufacturers use different technologies (linear stepper motor, encoder, etc.) to get the most accurate repositioning. Our measurements mainly involved linear 2-phases step motor stages, as these are the most popular on microscope systems. Most measured metrics were within the manufacturers’ specifications. We define as limit values the repeatability value given by the manufacturer.

As repeatability gets worse when the distances between two positions increase, one should keep in mind that although the stages are within manufacturer’s specifications, repeatability doesn’t imply a perfect repositioning. For instance, we found that repositioning across 30 mm distances (e.g., distance between different wells of a multiwell plate) is less precise than values obtained across 3 or 0.3 mm distances. Notably, our measurements were performed considering the stage as part of the microscope and under ‘real’ conditions, which is not always in accordance with the manufacturers’ protocols and given values in the datasheet (the stage is considered as an isolated item).

### Camera noise

Characterizing a detector may prove quite complex, as several technologies are involved and because it is the last element of the light path of a microscope. Metrics for single-point detectors (APD, PMT, HyD) were proposed in the past ^5, 23^. We focused on array detectors (CCD, EMCCD and especially sCMOS cameras) and propose an easy protocol to measure read noise-related metrics, including dark offset and DSNU values that are useful for further image quantifications.

We defined a first VAR (%) metric as the ratio of the manufacturer’s specification to the measured read noise value. Most measured values were less than 10% variation from 100%, and we considered this value as our experimental tolerance value. When applicable, on-chip warm pixel (or defect or blemish) correction yielded lower noise values. Warm pixels (i.e., pixels showing significantly higher signal than the average pixels) are more frequent with sCMOS cameras. As conversion from charge to the digital output involves a single electronic chain in CCD/EM-CDD cameras, the noise is relatively uniform across the array. Charge-to-voltage conversion in sCMOS is performed at the pixel level, and each pixel column has independent amplifiers and analog-to-digital converters. Thus sCMOS sensors are prone to pixel-to-pixel variability (including read noise, dark noise and dark current variability). A hot pixel correction may compensate for this. For the sensor characterization, it is preferable to avoid warm pixel correction. It can hide some information on the number of warm pixels and their evolution in time (Figure 8b). For QC studies, one should always use the same mode when monitoring the read noise and the DSNU in time.

It should be noted that the number of warm pixels may increase with the exposure time due to the dark noise influence or the cosmic rays. Cameras are constantly subjected to cosmic rays, which implies an increase in the number of hot pixels blocked at the saturation values ^47–49^. Here we define a hot pixel as a pixel that does not respond to light.

Which noise needs to be taken into account, rms or median, is often debatable. Our protocol calculates both, and we represent them for the VAR metric in Figure 8d. According to theory, for CCD and EMCCD cameras, the rms and median model give similar read noise values, since the readout in these cameras is serial, and an identical read noise across the whole chip is expected. In sCMOS sensors, the noise distribution is not symmetrical. Thus both approximations, gaussian (rms) and median, are justified. Our interest here is to follow the calculated values over time and compare those with the ones in the camera datasheet. We recommend comparing with the value that is given in the datasheet and in any case contact the manufacturer to request the missing values. The median value provides information about the general magnitude of the noise. Together with the rms value, they indicate the spread of the read noise distribution.

Following the read noise over time can be done by calculating the STAB_noise_ metric. We found that this metric is always higher than 97%, which we consider an experimental tolerance value.

It should be mentioned that for a detector characterization, the linearity and the non-uniformity measurements of the photon response are of great importance. Of equal importance is the measurement of the sensitivity, which places the detector in a “detection system” frame, as the measured values can be influenced by the rest of the microscope components. These measurements require reference samples, nicely uniform illumination, and calibrated detectors to compare ^50^. They also require costly equipment and entail skill and time commitment levels that cannot be reasonably expected from most microscope users or facility staff. We thus do not recommend these protocols in that study.

## Conclusion

In this study, we used simple tools, as affordable as possible such as self-made bead slides, plastic fluorescent slides, and a power-meter. Regarding slide QC samples, many others exist, such as the Argolight slide, GATTAquant beads or nanorulers PSF Check slides, etc and would have also been an option. However, all the above solutions present advantages but also drawbacks (price or guaranteed performance for a limited duration in time) which influenced our choice of using the three tools mentioned above.

We described acquisition protocols that are simple, robust, reproducible, open-source, and do not take more than 10 min to acquire or at least to launch. Nevertheless, all these acquisitions require that the user is well trained to perform experiments in a correct and reproducible manner. In the future, the possibility of such protocols being performed with a single sample slide by software in an all automatized acquisition process would be an advantage, since the human factor can be skipped.

Every acquired data needs to be analyzed in a reproducible manner as well. To allow this, we developed and proposed the usage of the ImageJ/Fiji plugin MetroloJ_QC. For quality monitoring, we performed many acquisitions over time and proposed several metrics and limit values. When possible, we tried to compare the measured values with the values given by the theory, such as the resolution for PSF, the co-registration or even the mechanical drift. For some other cases we referred to mechanical specifications given by constructors as for the stage repositioning or the read noise for detectors. We also tried as much as we could, to choose the limit value relevant for the experiments in biology, what we call “Lab practical” in Figure 9.

**Figure 9:**
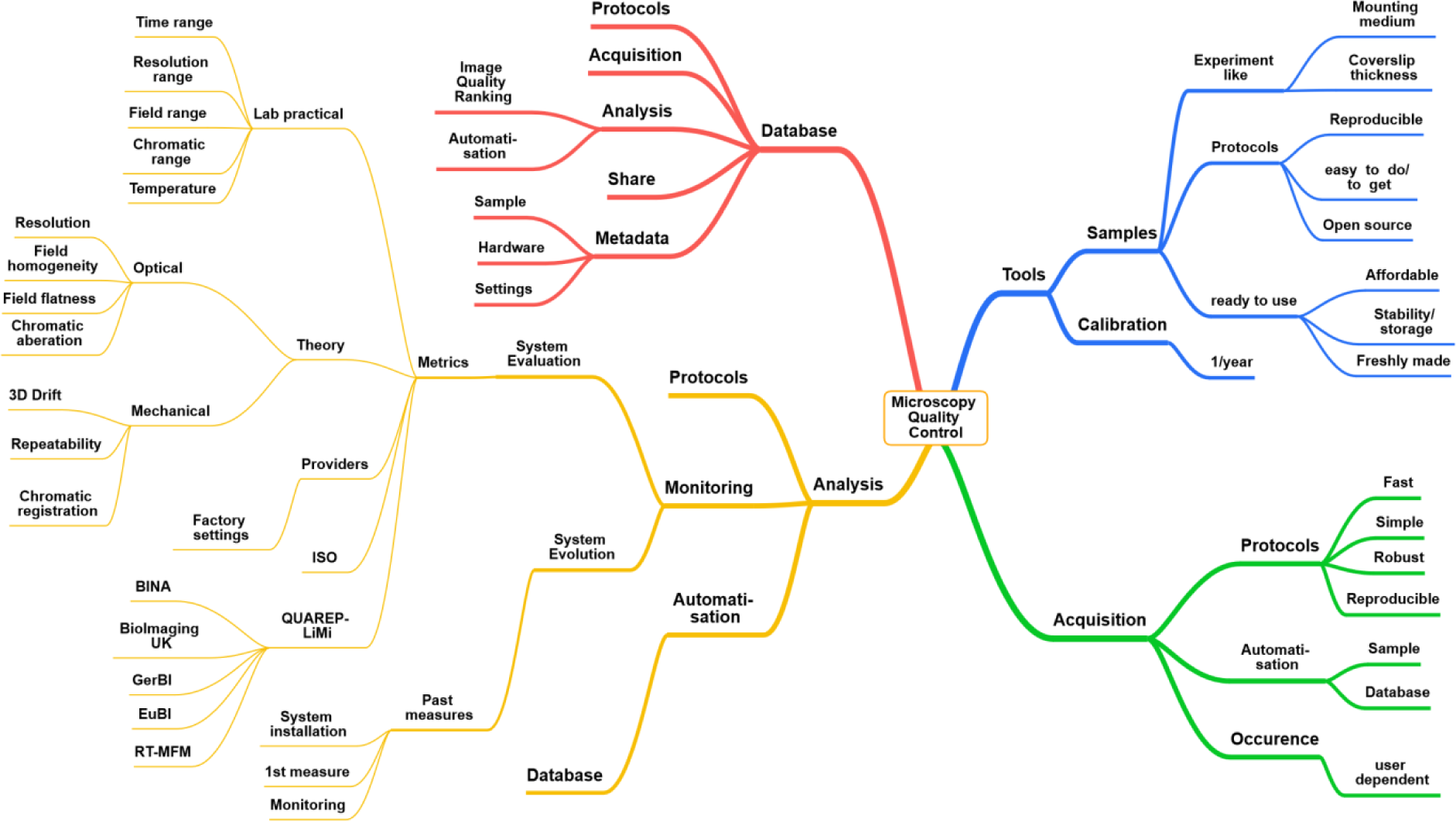
Heuristic map of Quality Control in light microscopy

Figure 9 shows in a heuristic way an idea of what Quality Control can be for fluorescence microscopy. We describe the essential aspects of Acquisition and Analysis Tools, Acquisition protocols and find some experimental tolerance values (Table 2).

**Table 2:** The basic metrics for monitoring the performance of a light microscope over time with the experimental limit values and the main trouble shooting cases.

| Monitoring | Metrics | Limit values | Trouble shooting |
| --- | --- | --- | --- |
| Point Spread Function | FWHM XY exp/theory | <1.5 | Optical aberrations, dirty or damaged objective, correction collar, excitation light alignment and polarization, pinhole alignment |
|  | FWHM Z exp/theory | <2 |  |
|  | Lateral asymmetry ratio LAR | Case dependent |  |
| Field Illumination | Uniformity U | >40% | Camera area, beam size at the back focal plane of the objective, dirty or damaged optics compounds |
|  | Centering C | > 50% | UV corrections, source and system alignment |
| Co-registration | Ratio $r_{\text{exp}}/r_{\text{ref}}$ | 1 | Chromatic aberrations, objective cleaning, dichroic positions in filter cubes (WF microscopy) |
| Illumination power stability | Stability $\text{STAB}_{\text{power}}$ | >97% | laser dying, defective optical components (AOTF, AOM, fiber) |
| | Standard Deviation $\text{STD}_{\text{normalized intensity}}$ | <0.2 | |
| Stage drift | Mean velocity after stabilization | <0.05 $\mu\text{m}/\text{min}$ | Thermal variations, air-flow, humidity, mechanical instability |
| | Mean velocity before stabilization | <0.1 $\mu\text{m}/\text{min}$ | |
| | $\tau_{\text{stabilization}}$ | <120 min | |
| Stage positioning repeatability | $\sigma_{\text{positioning deviation}}$ | Stage specifications | Thermal variations, air-flow, humidity, mechanical instability, stage drift |
| Camera read noise | VAR $\text{STAB}_{\text{noise}}$ | <10%<br>>97% | Electronics issue, sensor temperature, cosmic rays, camera and/or image sensor aging |

We consider that these metrics complement the ISO norm^13^ and hopefully will be further enriched by the QUAREP-LiMi, an international initiative started in April 2020 and whose role is to establish guidelines for quality assessment in light microscopy ^51^.

These metrics have to be monitored over time, first in a simple table. However, ideally, an image data management database would be the best solution to process the analysis and obtain the metrics as quality control images are imported ^52–54^. Furthermore, this database would be able to integrate these values in the metadata of the acquired images^55^. In the end, these metrics would then allow an open and accessible inspection of the quality of the used microscope system of every single image and would be a key contribution to an automatic image quality ranking assessment.

## Acknowledgements

The authors would like to thank Sebastian Beer and Bruno Emica from Hamamatsu Photonics and Gerhard Holst from PCO for their valuable support and discussion on the metrics of the QC of detectors. We would like to thank Kees van der Oord (Nikon), Christian Schulz (Leica Microsystems) and Ralf Wolleschensky (Zeiss) for their helpful input on PSF issues. We also thank Arnd Rühl (Märzhäuser) for his invaluable advice on the stages part of this manuscript. We are deeply indepted to Ulrike Boehm for her critical reading of the manuscript. We finally thank the french RTmfm community for the fruitful exchanges throughout the past years. The Montpellier Ressources Imagerie (BCM), PICT (Institut Curie), Bordeaux Imaging Center and IMAG’IC (Institut Cochin) are part of the National Infrastructure France-BioImaging supported by the French National Research Agency (ANR-10-INBS-04). The work was supported by the french technological network RTmfm of the CNRS « Mission pour les Initiatives Transverses et Interdisciplinaires ».

