## Supplemental Figures for "Long-term quality assessment and monitoring of light microscope performance through accessible and reliable protocols, tools and metrics"

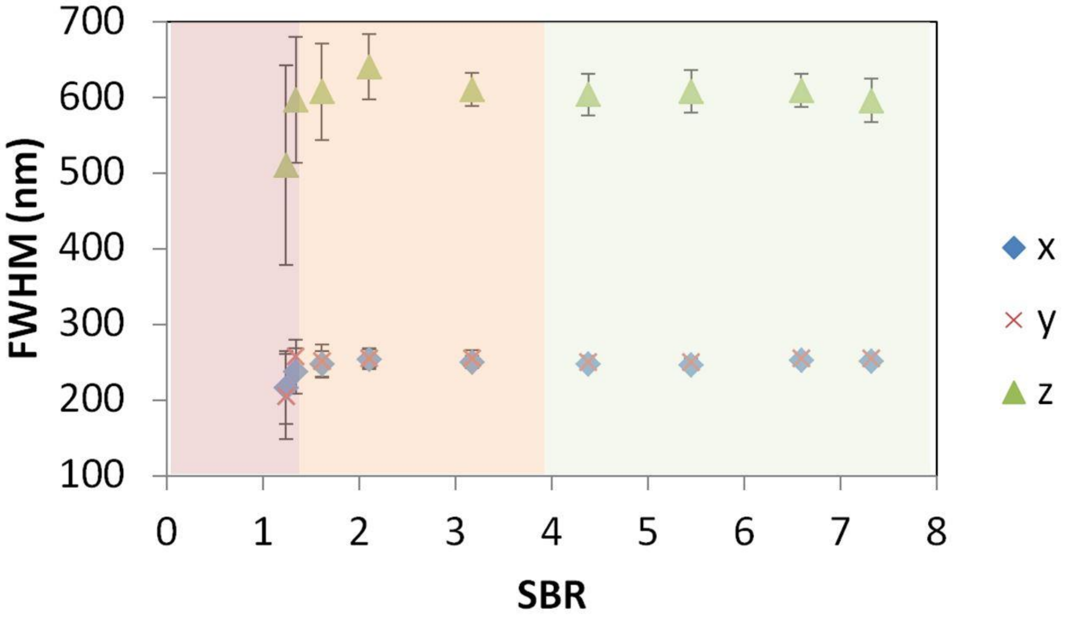


**Supplemental Figure 1: Effect of the S/B ratio on FWHM estimation precision for wide-field microscope.** The same three field of views of 29 beads were acquired for different exposure times and excitation intensities (with an upright Zeiss microscope, at the GFP channel, 63x lens, NA 1,4). The S/B ratio is estimated using the segmented bead mean intensity value (Signal) and the mean value of a 1um thick background annulus around the segmented bead (Background). As S/B is increasing, the FWHM estimation accuracy gets better. For very low S/B ratio the FWHM values decrease. We distinguish three tolerance zones. The green zone is the one presenting the lowest errors and the best fit (R2 equal to 1). The orange zone presents a higher error and a worse fit (0,93<R2<0,99). The red zone presents both high error and bad fit (R2 < 0,9).

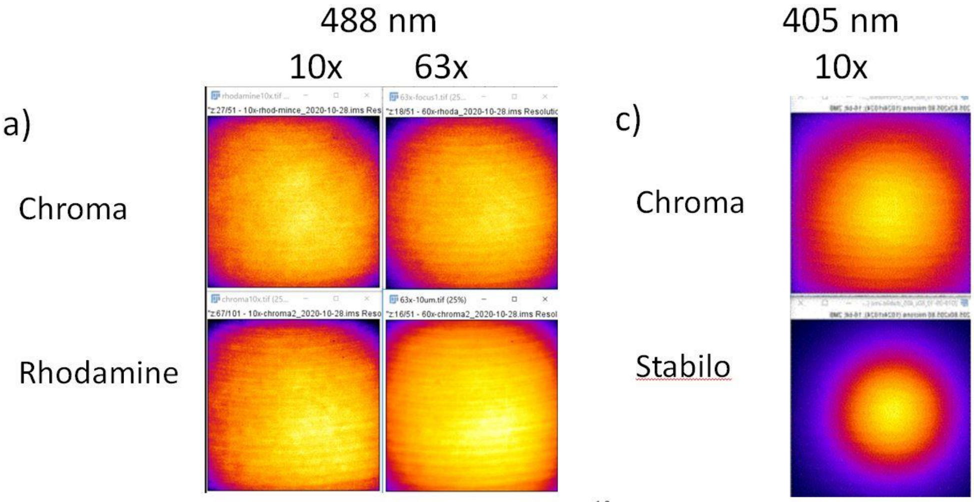


b)

| | Centering | Uniformity |
|---|---|---|
| Chroma focus | **59** | **63** |
| Chroma 10um | 57 | 73 |
| Chroma 30um | 60 | 64 |
| Rhodamine | **55** | **78** |

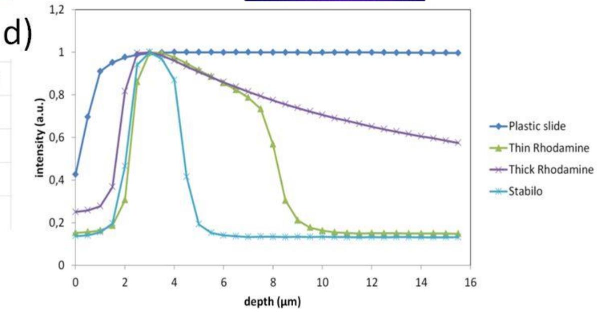


**Supplemental Figure 2: Field illumination (spinning disk microscope) using either a green chroma plastic slide or a glass coverslide/coverslip configuration with a rhodamine layer/green stabilo mark.** a) Field illumination images at 488nm excitation for a 10x and 63x with a plastic and rhodamine slide. b) C and U metrics for a). Acquisition was set at the plastic/glass or rhodamine/coverslip interface, or 10/30µm deep within the plastic/rhodamine layer. c) comparison of field illumination images (405nm excitation, a 10x lens) using either a plastic slide or a stabile mark. d) Effect of focal plane depth on emission intensity using either a plastic slide, a stabile mark or thin/thick layers of fluorescent rhodamine dye (63x objective, ex.488nm).

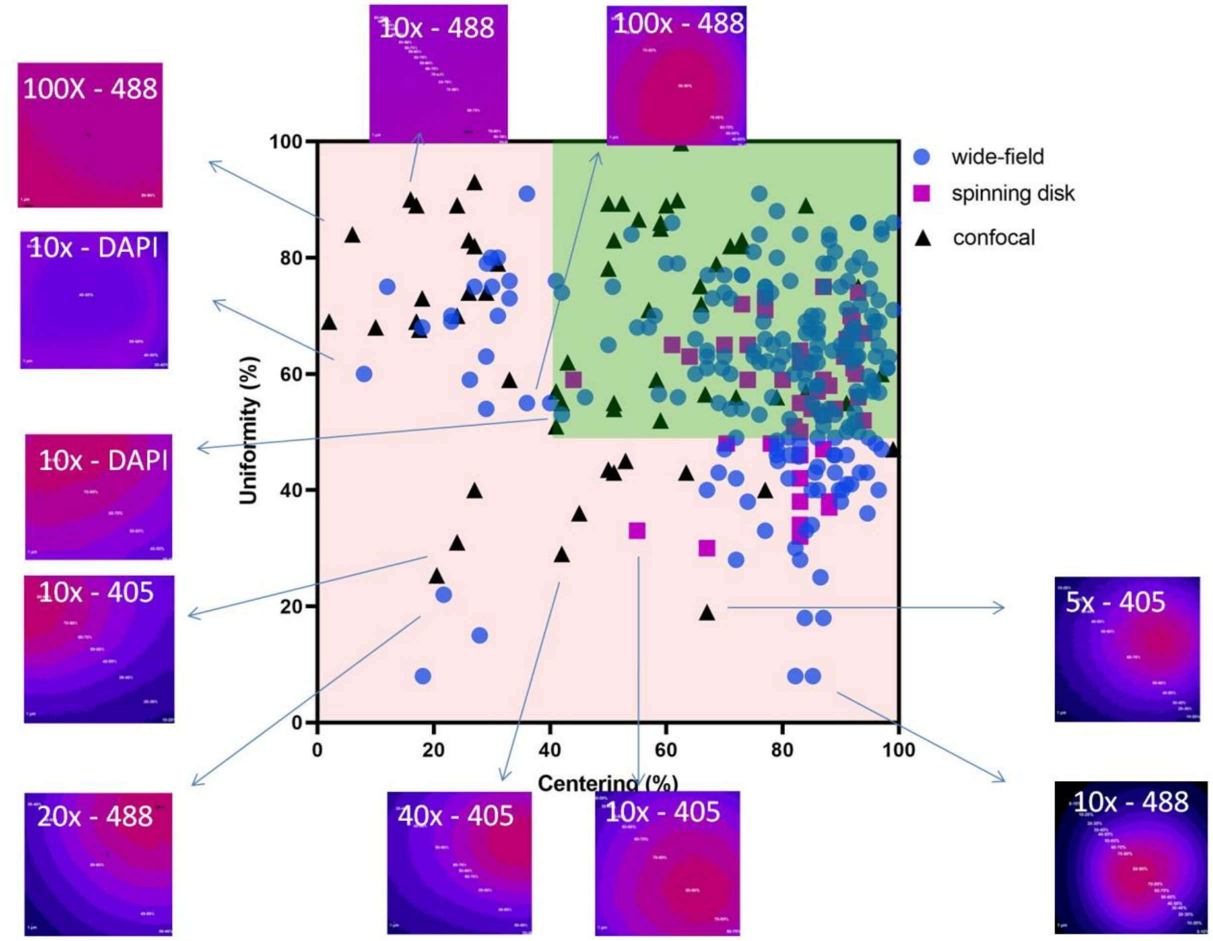


**Supplemental Figure 3: Distribution of C and U.** Detailed version of Figure 3, showing the extreme cases for both metrics.

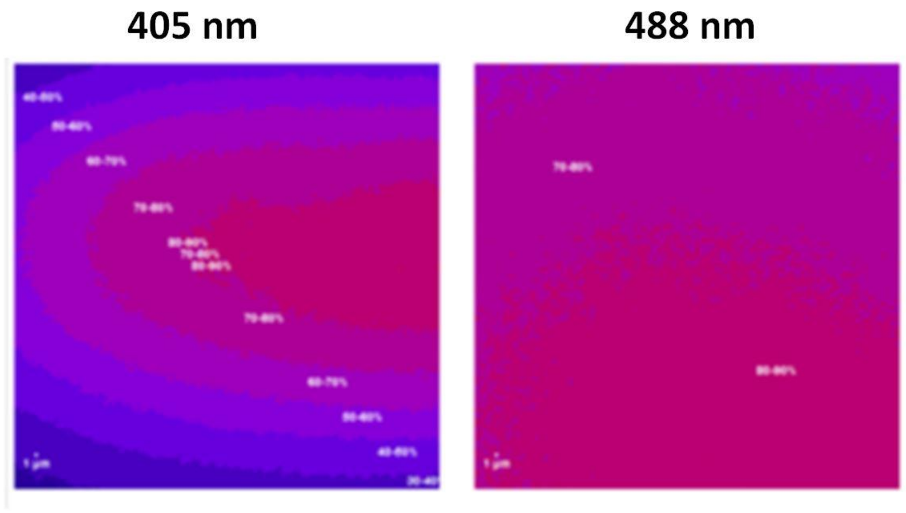


| | 405 nm | 488 nm |
|---|---|---|
| Centering % | 45 | **27** |
| Uniformity % | **36** | 82 |

**Supplemental Figure 4: Field illumination metrics of a LSM 880 confocal microscopefor a 63x 1,4 NA lens for 405 and 488 laser excitation.**

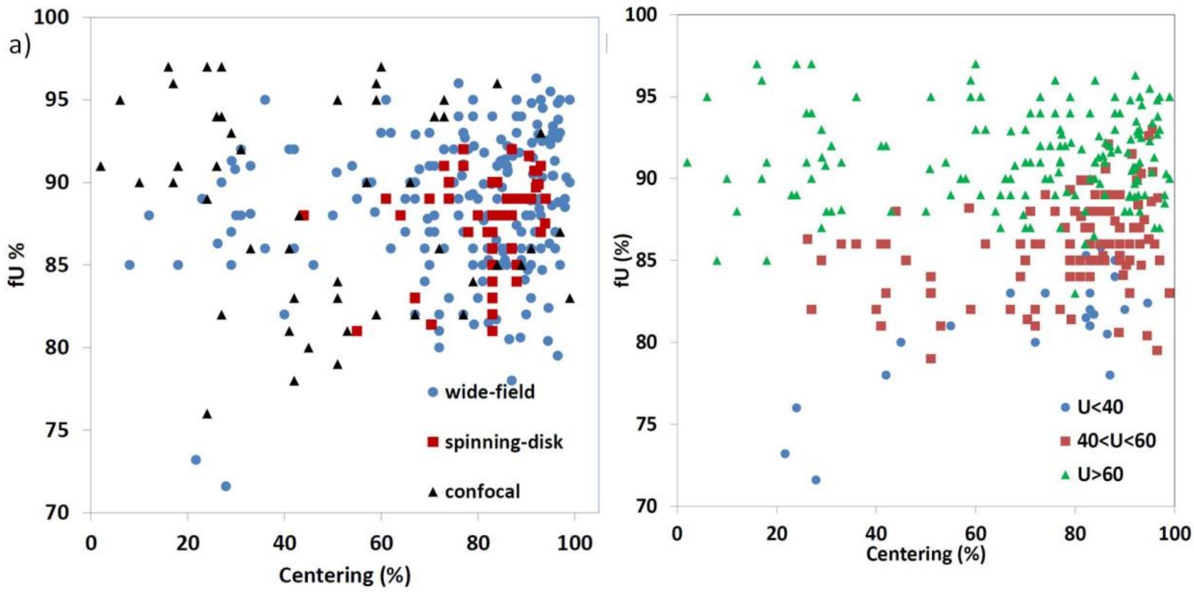


**Supplemental Figure 5: Field illumination flatness fU and Centering metrics.** a) fU vs Centering for the 3 microscopy techniques of wide-field, spinning disk and confocal. b) fU vs Centering for all 3 microscope techniques, divided in 3 groups depending on the Uniformity value.

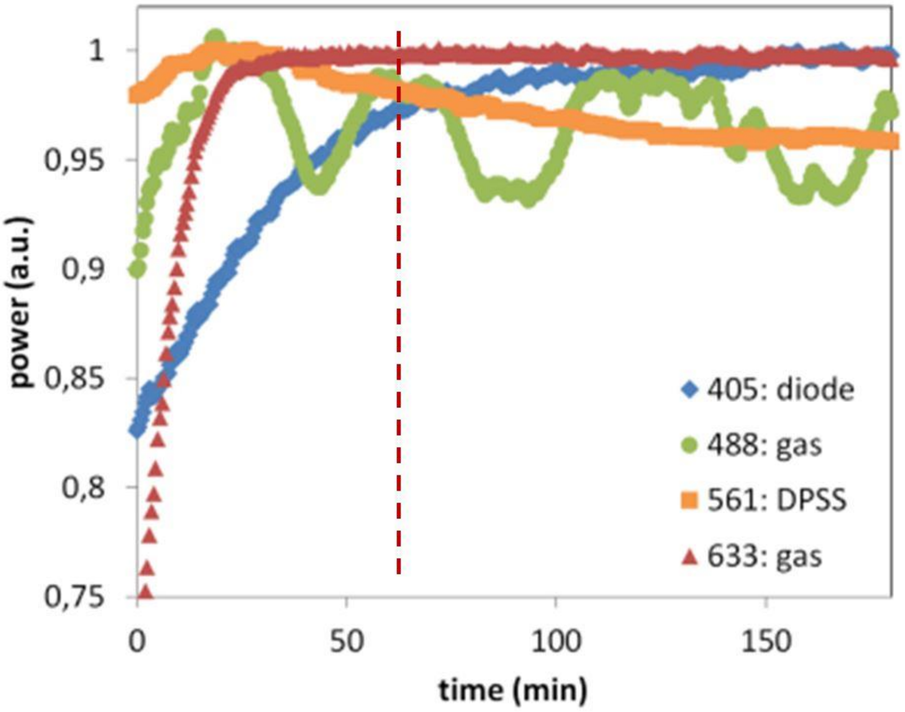


**Supplemental Figure 6: Study to find the warming up time of lasers.** Stability of 4 different laser types on a LSM microscope just after switching each source on. Red dashed line points out the average 60 min warming up time for all sources, taken into account in Figure 5.

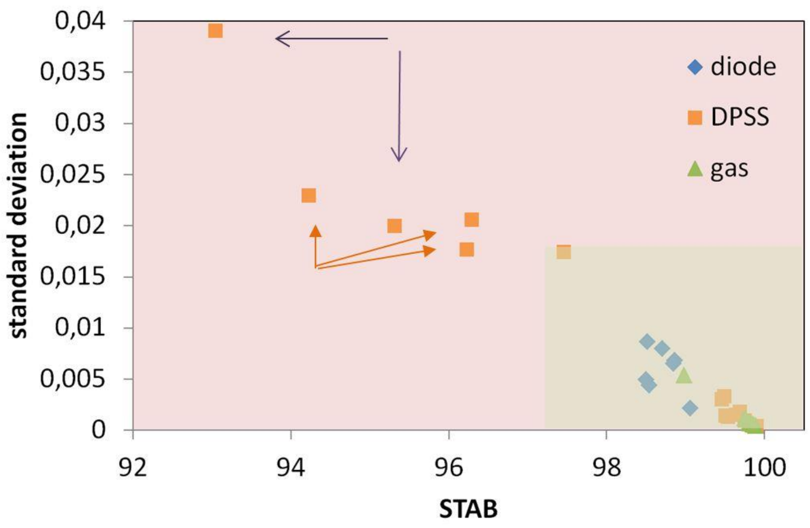


**Supplemental Figure 7: Illumination power short-term stability.** Laser power stability STAB vs standard deviation of normalized to 1 laser power values over 5min. With low stability, the lasers come from the same microscope. The magenta arrows point 2 DPSS lasers of the same microscope pointed out with red arrow in Figure 5d. The orange closed arrows point 3 DPSS lasers are coming from the same microscope.

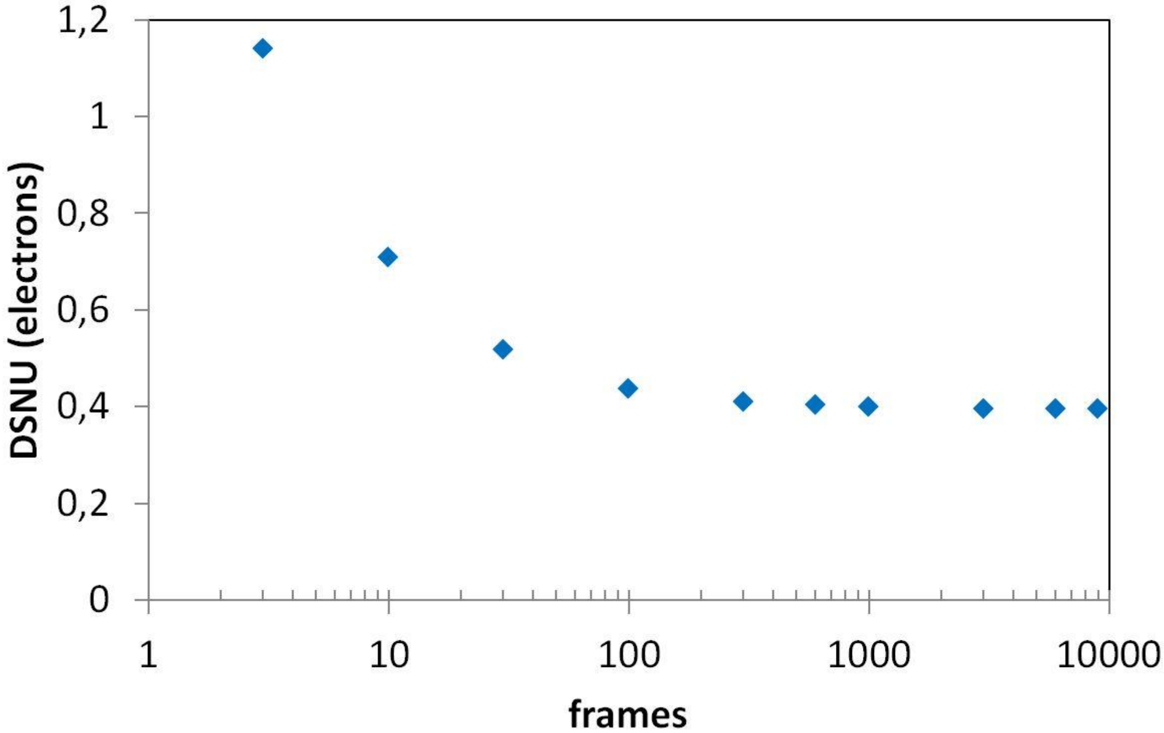


**Supplemental Figure 8 : Distribution of DSNU values calculated from different dark image dataset sizes.** Images were acquired using an Andor Zyla 4.2 sCMOS camera. Using a 100 frames dataset, the calculated DSNU is 0,438 e-, while for 9000 frames the DNSU is 0,395 e-.
